# Decrowding Expansion Pathology: Unmasking Previously Invisible Nanostructures and Cells in Intact Human Brain Pathology Specimens

**DOI:** 10.1101/2021.12.05.471271

**Authors:** Pablo A. Valdes, Chih-Chieh (Jay) Yu, Jenna Aronson, Yongxin Zhao, Joshua D. Bernstock, Deepak Bhere, Bobae An, Mariano S. Viapiano, Khalid Shah, E. Antonio Chiocca, Edward S. Boyden

## Abstract

Proteins are densely packed in cells and tissues, where they form complex nanostructures. Expansion microscopy (ExM) variants have been used to separate proteins from each other in preserved biospecimens, improving antibody access to epitopes. Here we present an ExM variant, decrowding expansion pathology (dExPath), which can expand proteins away from each other in human brain pathology specimens, including formalin-fixed paraffin-embedded (FFPE) clinical specimens. Immunostaining of dExPath-expanded specimens reveals, with nanoscale precision, previously unobserved cellular structures, as well as more continuous patterns of staining. This enhanced molecular staining results in observation of previously invisible disease marker-positive cell populations in human glioma specimens, with potential implications for tumor aggressiveness. dExPath results in improved fluorescence signals even as it eliminates lipofuscin-associated autofluorescence. Thus, this form of expansion-mediated protein decrowding may, through improved epitope access for antibodies, render immunohistochemistry more powerful in clinical science and diagnosis.

## Introduction

Immunohistochemistry, a technique that has revealed fundamental insights in biology and is applied in diverse clinical settings, relies on the ability of antibodies to access epitopes on proteins embedded in intact cells and tissues. The most commonly used antibodies are IgG-class immunoglobulins, which have a non-negligible size of 14.5 x 8.5 x 4.0 nm^1,2^. Due to this non-negligible size, target epitopes in fixed tissues are often physically inaccessible to antibodies^3–14^.

Expansion microscopy (ExM) enables physical expansion of biological specimens, thereby permitting nanoscale resolution imaging on diffraction-limited microscopes^15,16^. Briefly, ExM starts by covalently anchoring biomolecules, or labels against targeted biomolecules, to a swellable hydrogel network densely and evenly synthesized throughout a preserved biological specimen. Then, an enzymatic or protein-denaturing treatment softens the mechanical properties of the specimen. Water then causes the polymer network to expand, and thus the anchored molecules to be pulled uniformly away from one another. Given the difficulty of labeling many epitopes in their natural, densely packed state, we asked whether, in human tissues of interest in pathology and medicine, conventional antibodies introduced in the post-expansion, i.e. decrowded, state could access previously undetectable epitopes.

Some expansion protocols have been shown to be capable of preserving protein antigens throughout the expansion process (**Supp. Table 1**)^17–25^, and are thus compatible with post-expansion immunostaining. However, most of these existing post-expansion staining protocols either required specialized fixative compositions^17,18,21,22,24^, and thus are incompatible with archival clinical samples, or they showed incomplete softening with tissue cracks and anisotropy^19^, or had uncharacterized nanoscale isotropy^20^. In addition, none of these studies underwent quantitative comparison of structures or cells in the same specimen of human tissue compared with pre- versus post-expansion staining, key to understanding whether the decrowding of proteins contributed to visualization of previously invisible structures.

We previously developed expansion pathology (ExPath), a form of ExM that prepares human specimens preserved through various standard fixation and archival protocols, for expansion microscopy, using pre-expansion antibody staining to provide molecular contrast^6^. Here we present decrowding ExPath (dExPath), an expansion pathology variant that preserves protein epitopes for post-expansion staining, while still expanding human tissues isotropically. dExPath can be applied to formalin-fixed paraffin-embedded (FFPE) human clinical tissues, as well as other standard formats of interest in basic and applied biology (e.g., 4%-paraformaldehyde (PFA)-fixed mouse brain tissue). We validated dExPath systematically, comparing, within the same specimen of human brain tissue, immunostaining intensity and continuity between pre- and post-expansion staining, showing improvements in both intensity and continuity, and even revealing entirely new features, and new disease marker-bearing cell populations (in human glioma specimens), that were previously invisible. Furthermore, dExPath specifically eliminates the autofluorescence associated with lipofuscin, an aggregated waste product commonly found in brain tissue, beyond just the tissue-wide autofluorescence reduction resulting from the loss and dilution of autofluorescent molecules in prior expansion protocols^6^. dExPath also supports multi-round immunostaining, enabling highly multiplexed (here demonstrated with 10 stains, but supporting likely far more) imaging of protein targets within the same human brain specimen. We anticipate dExPath to open up many new experimental capabilities in the study of detailed protein assemblies and cellular structures in brain specimens, and perhaps other tissue types as well.

## Results

### Rationale for the dExPath technology

We first prepared tissue to enter the expansion pipeline (**Fig. 1A**; e.g., involving tissue deparaffinization and re-hydration, for FFPE samples)^6^, followed by protein anchoring and gel formation (**Fig. 1B**). In contrast to the original ExPath protocol, which uses a strong protease digestion to soften the specimen (feasible because fluorescent antibodies, which are partly protease-resistant, are applied pre-expansion and directly anchored to the polymer network for later imaging), we here created a buffer to maximally enable protein separation for post-expansion staining. We used higher levels of sodium dodecyl sulfate (SDS) (20% weight/volume (w/v)) than in earlier protein-preserving protocols (proExM, 1%; mExM, 4%; MAP/U-ExM/miriEx/pan-ExM/ExR, 5.8%; for other protein-preserving protocols, see **Supp. Table 1**)^17–24^, reasoning that this could help with isotropic expansion by better converting proteins to a denatured state, and minimizing non-covalent intra- and inter-protein interactions that could potentially hinder molecular separation and tissue expansion^26,27^. We also included a new ingredient, β mercaptoethanol (100 mM), a reducing agent, which we reasoned could help with isotropic expansion by cleaving inter-molecular disulfide bridges between structural components of the tissue^25–30^. We also used the same high level of ethylenediaminetetraacetic acid (EDTA) (25 mM) as in the original ExPath protocol, which showed this to be useful for isotropic tissue expansion^6^, possibly through de-stabilization of metal-mediated protein interactions in the extracellular matrix (ECM)^27–29^. We used a higher temperature than in the original ExPath protocol, as used in a form of proExM that uses autoclaving to expose samples to 121°C **(Fig. 1C**) to strongly denature, and loosen disulfide bonds and fixation crosslinks between proteins in the sample, allowing them to separate from one another during subsequent washes (which drives partial tissue expansion, i.e., ∼2.3x; **Fig. 1D**). Antibodies are applied at the post-decrowding state (**Fig. 1E;** see **Supp. Table 2** for antibodies used in this work); staining is performed at the partially expanded [∼2.3x] state, instead of the fully expanded [∼4x] state, because full expansion requires sample immersion in deionized water, a low-ionic-strength environment that hinders antibody binding, consistent with earlier post-expansion immunostaining protocols^17–22^. Multiplexing is possible because these antibodies can be stripped using the same buffer, and then new antibodies applied (**Supp. Fig. 1**), a strategy previously demonstrated by other post-expansion staining protocols but not on human tissues^18,20,31^. High grade glioma tissues are known to undergo abnormal endothelial proliferation, leading to some areas of tissue with abnormally large amounts of vascularity and extracellular matrix (ECM). These specific areas in tissue samples can be identified under conventional diffraction limited clinical microscopy staining (e.g., hematoxylin and eosin staining)^32^. These areas can present a challenge to isotropic expansion of tissue using dExPath^6^. To address this additional challenge in this specific and easily identified pathological state, we devised a modified form of the dExPath protocol using collagenase treatment prior to softening (**Supp. Fig. 2**). In summary, our dExPath protocol was designed to provide a methodology for isotropic tissue expansion, enabling preservation, and post-expansion as well as multiplexed staining, of decrowded proteins in both normal and pathologic human and rodent brain tissues.

**Fig. 1.**
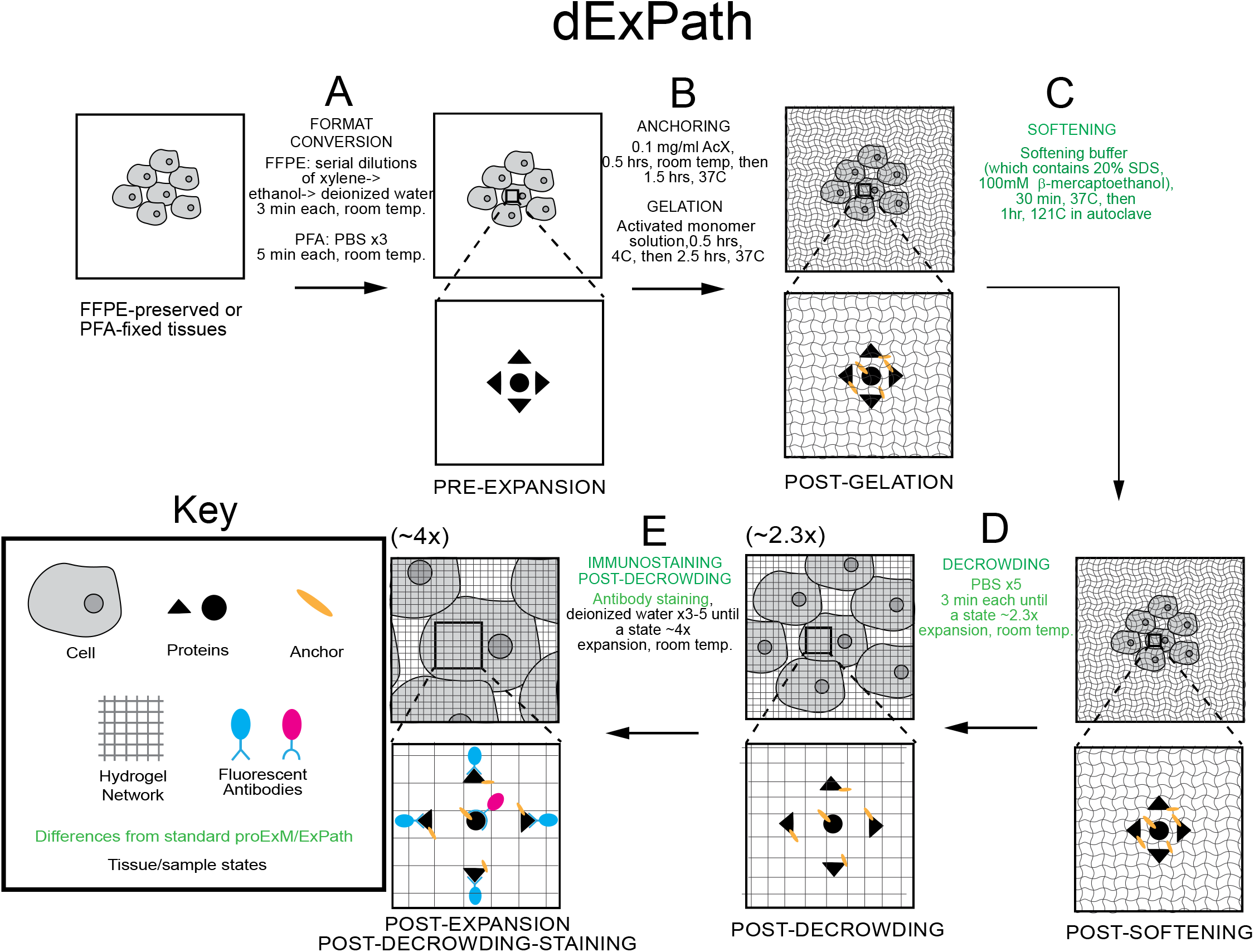
Decrowding expansion pathology (dExPath) for post-expansion immunostaining of human tissue and other formaldehyde-fixed specimens. (A-E) Workflow for expanding formalin-fixed paraffin-embedded (FFPE), or formaldehyde-fixed, human (or mouse) brain specimens, enabling post-decrowding immunostaining. Key modifications of published proExM and ExPath protocols are shown in green. PFA, paraformaldehyde; PBS, phosphate buffered saline; RT, room temperature; AcX, Acryloyl-X; SDS, sodium dodecyl sulfate. For steps after decrowding (D), linear expansion factor of the hydrogel-specimen composite is shown in parentheses above the schematic of the step. (A) Tissue samples undergo conversion into a state compatible with expansion. (B) Tissue samples are treated so that gel-anchorable groups are attached to proteins, then the sample is permeated with an expandable polyacrylate hydrogel. (C) Samples are incubated in a softening buffer to denature, and loosen disulfide bonds and fixation crosslinks between, proteins in the sample. (D) Softened samples are washed in a buffer to partially expand them. (E) Samples are stained and then expanded fully by immersion in water.

### Validation of dExPath expansion isotropy in brain tissue

We validated the isotropy of dExPath on normal and diseased FFPE-preserved, 5-µm-thick brain tissues (a standard thickness for clinical samples), using the same pre-vs-post distortion analysis used for earlier expansion protocols^6,15,18,19,33,34^. We performed antigen retrieval followed by pre-expansion immunostaining against microtubule-associated protein 2 (MAP2, a neuronal dendritic marker)^35^, and the intermediate filament protein vimentin^36–38^, on normal human hippocampus (**Fig. 2A**) and on high-grade glioma tissues (located in the human cortex or white matter) (**Fig. 2B**), respectively. We applied a workflow for immunostaining FFPE-preserved clinical tissues^6,39–41^ to obtain pre-expansion images of tissues using a super-resolution structured illumination microscope (SR-SIM) (**Fig. 2A-B**). Next, we performed dExPath (but, using pre-expansion staining prior to anchoring and gelation, using a protocol modified to facilitate distortion comparison between pre- and post-expansion images of the same sample, outlined in **Supp. Fig. 3**) on the immunostained tissues, obtaining post-expansion images of the same fields of view of the same samples (**Fig. 2C-D**) using a confocal microscope. We observed low distortion between pre- and post-expansion images of the same fields of view, similar to previous versions of ExM applied to mouse brain tissue^6,15,19^ **Fig. 2E**; ∼4% root mean squared (RMS) error over distances of ∼10 µm; n = 4 samples, each from a different patient; **Fig. 2F**; ∼3% RMS error over distances of ∼10 µm; n = 3 samples, each from a different patient). In specific instances of high-grade glioma tissues with large amounts of ECM identified under conventional clinical microscopy, our modified form of dExPath using collagenase treatment prior to softening was used to compare pre- and post-expansion images of the same specimen, outlined in **Supp. Fig. 4**. We found similar results on high-grade glioma tissues with a high degree of extracellular matrix using collagenase treatment prior to softening (**Supp. Fig. 5**). Thus, dExPath isotropically expands archival clinical samples of FFPE normal brain and brain tumor tissues by ∼4x without the need for enzymatic epitope destruction^6,19^, or specialized fixatives^17,18,21,22^.

**Fig. 2.**
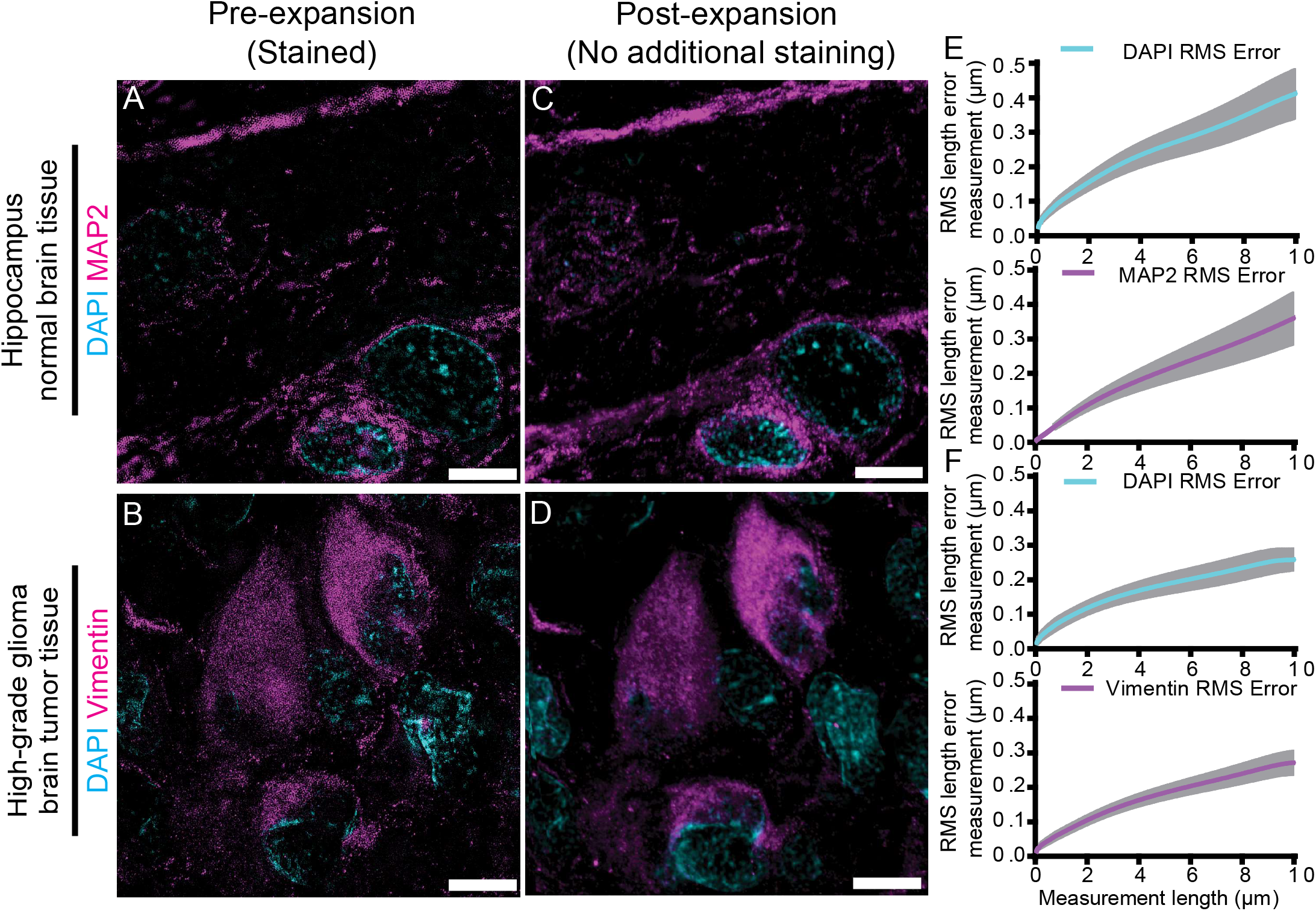
Isotropy of dExPath. (A-B) Representative pre-expansion super resolution structured illumination microscopy (SR-SIM) images of FFPE 5-µm-thick slices of normal human hippocampus (A, n = 4 samples, each from a different patient) and human high-grade glioma brain tumor tissue (B, n = 3 samples, each from a different patient) which underwent processing as in **Supp. Fig. 3A** (tissue deparaffinization, rehydration, antigen retrieval, and immunostaining), with immunostaining being for MAP2 and staining for DAPI (A), or staining for vimentin and DAPI (B). (C-D) Post-expansion images of the same fields of view as shown in (A-B), respectively. Specifically, samples underwent anchoring and gelation (as in **Supp. Fig. 3B**), softening (as in **Supp. Fig. 3C**), another round of DAPI staining, ∼4x linear expansion (as in **Supp. Fig. 3D**), and imaging with confocal microscopy. (E-F) Root-mean-square (RMS) length measurement errors obtained by comparing pre- and post-expansion images such as shown in A-D (n = 4 samples, each from a different patient, E; n = 3 samples, each from a different patient, F). Line, mean; shaded area, standard deviation. All images are sum intensity z-projections, either of SR-SIM image stacks (A-B), or confocal image stacks (C-D), both covering an equivalent tissue depth in biological units. Brightness and contrast settings: first set by the ImageJ auto-scaling function, and then manually adjusted (by raising the minimum-intensity threshold and lowering the maximum-intensity threshold) to improve contrast for the stained structures of interest but quantitative analysis in (E-F) was conducted on raw image data. Scale bars (in biological units: physical sizes of expanded samples divided by their expansion factors, used throughout this manuscript, unless otherwise noted): (A-D) 5 µm. Linear expansion factors: (C-D) 4.0x.

### dExPath removes lipofuscin autofluorescence, improving visualization of intracellular structures

Fluorescence microscopy of clinical tissues is often hindered by lipofuscin^42–49^, an autofluorescent (throughout the visible optical spectrum) waste material that is composed of aggregates of oxidized proteins, lipids, and metal cations, and that accumulates in many cell and tissue types^50–53^. We imaged regions with lipofuscin in normal human cortex (age: 19 – 45 years old), in the pre-expansion state (**Fig. 3A-D**) and in the post-expansion state (**Fig. 3E-H**), under 3 common fluorescent filter settings (488 nm excitation (abbreviated as “ex”)/525 nm emission (abbreviated as “em”); 561ex/607em; 640ex/685em), finding that lipofuscin fluorescence was an order of magnitude, or more, than background fluorescence (**Fig. 3D**; lipofuscin vs background: 488ex/525em, p = 0.00001; 561ex/607em, p = 0.00002; 640ex/685em, p = 0.00002; 2-tailed paired t-test; all t-tests were non-Bonferroni corrected; n = 4 tissue samples, each from a different patient). After dExPath, the autofluorescence from the lipofuscin was reduced to a level that was indistinguishable from background (**Fig. 3H**; lipofuscin vs background: 488ex/525em, p = 0.11; 561ex/607em, p = 0.07; 640ex/685em, p = 0.29; 2-tailed paired t-test; n = 4 tissue samples, each from a different patient). Classical ExPath still showed some lipofuscin autofluorescence post-expansion (**Supp. Fig. 6**). Using dExPath, structures that were previously masked by lipofuscin became detectable. Comparing the same location in the same specimen pre- and post-expansion, with stains against MAP2^35^, giantin (a Golgi-apparatus marker)^54,55^, and synaptophysin (a pre-synaptic marker)^56^ (**Fig. 3I-K**), some giantin staining overlapped with lipofuscin (compare **Fig. 3B** vs. **3J,** respectively), thus could be obscured by autofluorescence from lipofuscin (note, these images were obtained with the same microscope settings). As another example, human hippocampal tissues that underwent pre-expansion immunostaining against MAP2 (in the 488ex/525em channel) and glial fibrillary acidic protein (GFAP, a marker of astrocytes^36,57,58^; in the 640ex/685em channel) showed false positive fluorescence in the GFAP channel in somata of MAP2-positive cells, due to lipofuscin presence there (**Fig. 3L**). In contrast, post-decrowding, such false positive GFAP staining no longer appeared in the somata (**Fig. 3M**), because the lipofuscin was removed; that region only showed MAP2-positivity, as expected. Thus, dExPath-mediated lipofuscin removal has the potential to greatly improve detection of subcellular fluorescent signals in human tissues.

**Fig. 3.**
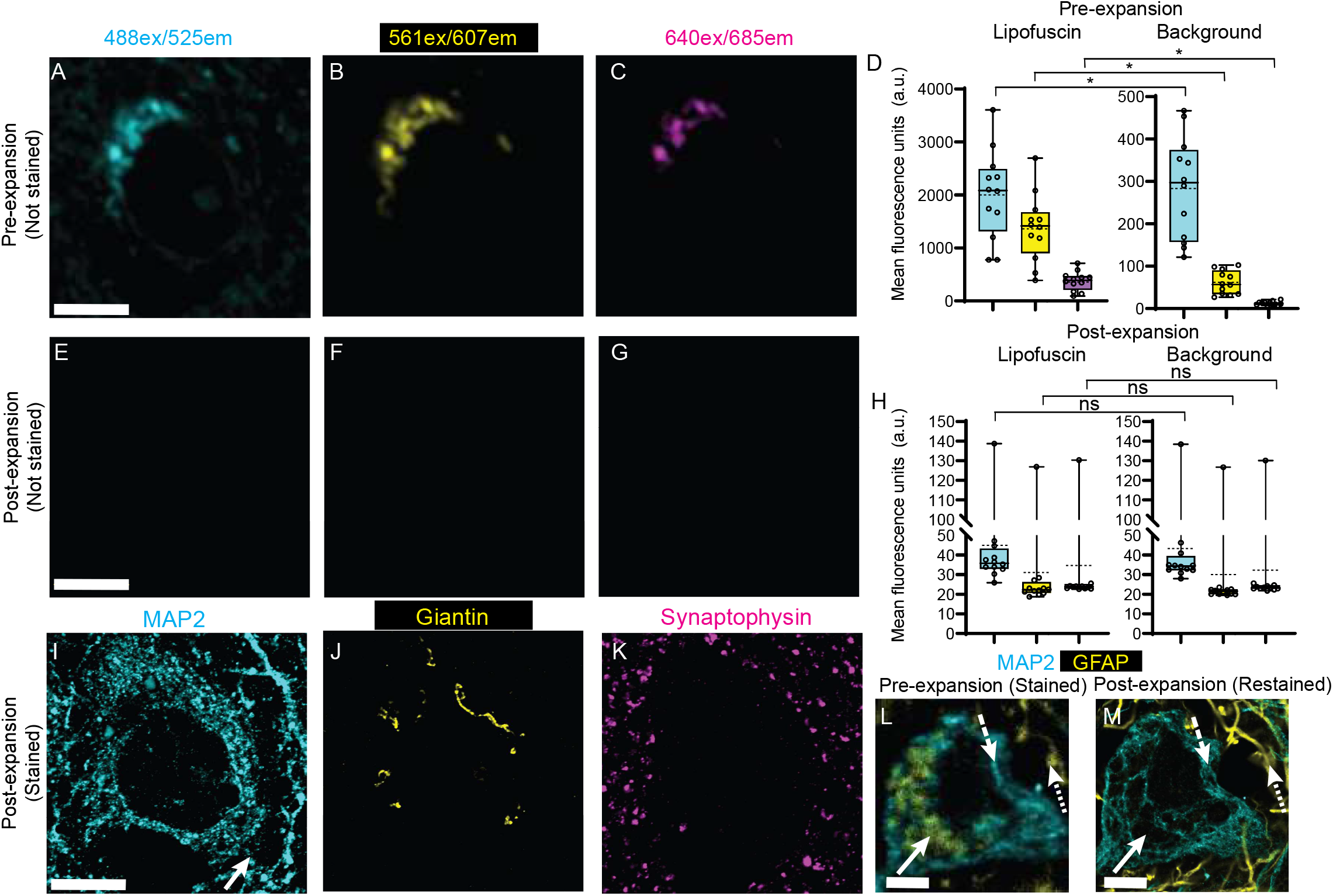
dExPath removal of lipofuscin autofluorescence. (A-C) Representative (n = 4 samples, each from a different patient) pre-expansion confocal images (single z slices) of a neuron of an FFPE 5-µm-thick sample of normal human cortex. The samples underwent format conversion (as in Fig. 1A, tissue deparaffinization and rehydration), and DAPI staining (images not shown in this figure; used for registration across images). Images were acquired for 3 common fluorescent filter settings: (A) a 488 nm excitation (abbreviated as “ex”) / 525 nm emission (abbreviated as “em”) channel; (B) a 561ex/607em channel; and (C) a 640ex/685em channel. (D) Mean fluorescence intensities from pre-expansion images, averaged across regions of interest (ROIs) that exhibited prominent lipofuscin (left bar graph), as well as across background ROIs (right bar graph); colors correspond to the colors of A-C (n = 4 tissue samples, each from a different patient). Brightness and contrast settings: first set by the ImageJ auto-scaling function, and then manually adjusted (by raising the minimum-intensity threshold and lowering the maximum-intensity threshold) to improve contrast for lipofuscin but quantitative analysis in (D) was conducted on raw image data. Box plot: individual values (open circles; 3 measurements were acquired from each patient), median (middle line), mean (dotted line), first and third quartiles (lower and upper box boundaries), lower and upper raw values (whiskers). Statistical testing: 2-tailed paired t-test (non-Bonferroni corrected) was applied to lipofuscin vs. background, for pre-expansion mean fluorescence intensities for each spectral channel. *, p < 0.05; ns, not significant. (E-G) Post-expansion confocal images after the sample from A-C was additionally treated with anchoring, gelation (as in Fig. 1B), softening (as in Fig. 1C), de-crowding (as in Fig. 1D), DAPI staining, and ∼4x linear expansion, without immunostaining at the post-decrowding state. Sum intensity z-projections of image stacks corresponding to the biological thickness of the original slice, taken under identical settings and of the same field of view as A-C and displayed under the same settings. (H) Mean fluorescence intensities, from post-expansion images, averaged across the same lipofuscin (left) and background (right) ROIs used in panel D, for the same samples as in panel D. Plots and statistics as in D. (I-K) Confocal images of the same field of view as in (E-G), but the sample was additionally immunostained, post-decrowding (as in Fig. 1E), for MAP2 (microtubule-associated protein 2), giantin, and synaptophysin (labeled with antibodies in the same spectral ranges as indicated above A-C), as well as stained for DAPI (not shown; again, used for alignment), and then re-expanded to ∼4x linear expansion; brightness and contrast settings: first set by the ImageJ auto-scaling function, and then manually adjusted (by raising the minimum-intensity threshold and lowering the maximum-intensity threshold) to improve contrast for stained structures. (L) Representative (n = 3 samples, each from a different patient) confocal image of a tissue sample of FFPE 5-µm-thick normal human hippocampus processed according to the protocol for pre-decrowding staining (for protocol schematic, see Supp. Fig 3A). Samples underwent format conversion, antigen retrieval, and pre-expansion immunostaining for MAP2 (488ex/525em channel) and GFAP (glial fibrillary acidic protein) (640ex/685em channel). Solid arrow indicates a region with lipofuscin aggregates (GFAP-like staining but found in a neuron); dashed arrow indicates MAP2 staining without lipofuscin; dotted arrow indicates GFAP staining. (M) Confocal image of the same field of view as (L). Tissues underwent softening and ∼4x expansion (protocol details in Supp. Fig. 3B-D), followed by decrowding, post-decrowding staining for MAP2 and GFAP, and expansion to ∼4x (as in Supp. Fig. 3E-F). Arrows, as in L. Brightness and contrast settings adjusted as in (I-K) to improve contrast for stained structures. Scale bars (in biological units): (A, E, I) 7 µm; (L, M) 5 µm. Linear expansion factors: (E-G, I-K) 4.3x; (M) 4.1x.

### dExPath enables visualization of decrowded proteins revealing previously invisible cells and structures

We next investigated whether post-expansion immunostaining could enable detection of previously inaccessible (that is, when in the conventional, crowded state) protein epitopes. To explore this possibility, we performed within-sample comparisons (i.e., pre- vs. post-expansion staining) using normal human hippocampus (**Fig. 4A-F**), supratentorial high-grade glioma tumor specimens (**Fig. 4G-R**), and low-grade glioma tumor specimens (**Fig. 4S-X**). Tissue samples were imaged pre-expansion, after antigen retrieval and antibody staining (**Fig. 4A, G, M, S**), as well as after expansion (**Fig. 4B, H, N, T;** in a state similar to ExPath in that antibodies were anchored prior to decrowding, but unlike ExPath, treated with a chemical (non-enzymatic) softening protocol that preserves epitopes); and after post-expansion re-staining with the same antibodies under the same conditions (**Fig. 4C, I, O, U;** experimental pipeline in **Supp. Fig. 3**). All tissue states were imaged using identical confocal imaging settings; we adjusted the raw images to generate those in **Fig. 4** by adjusting histograms (to the right of **Fig. 4A-C, 4G-I, 4M-O,** and **4S-U**) so that 1% of pixels were saturated in each picture.

**Fig. 4.**
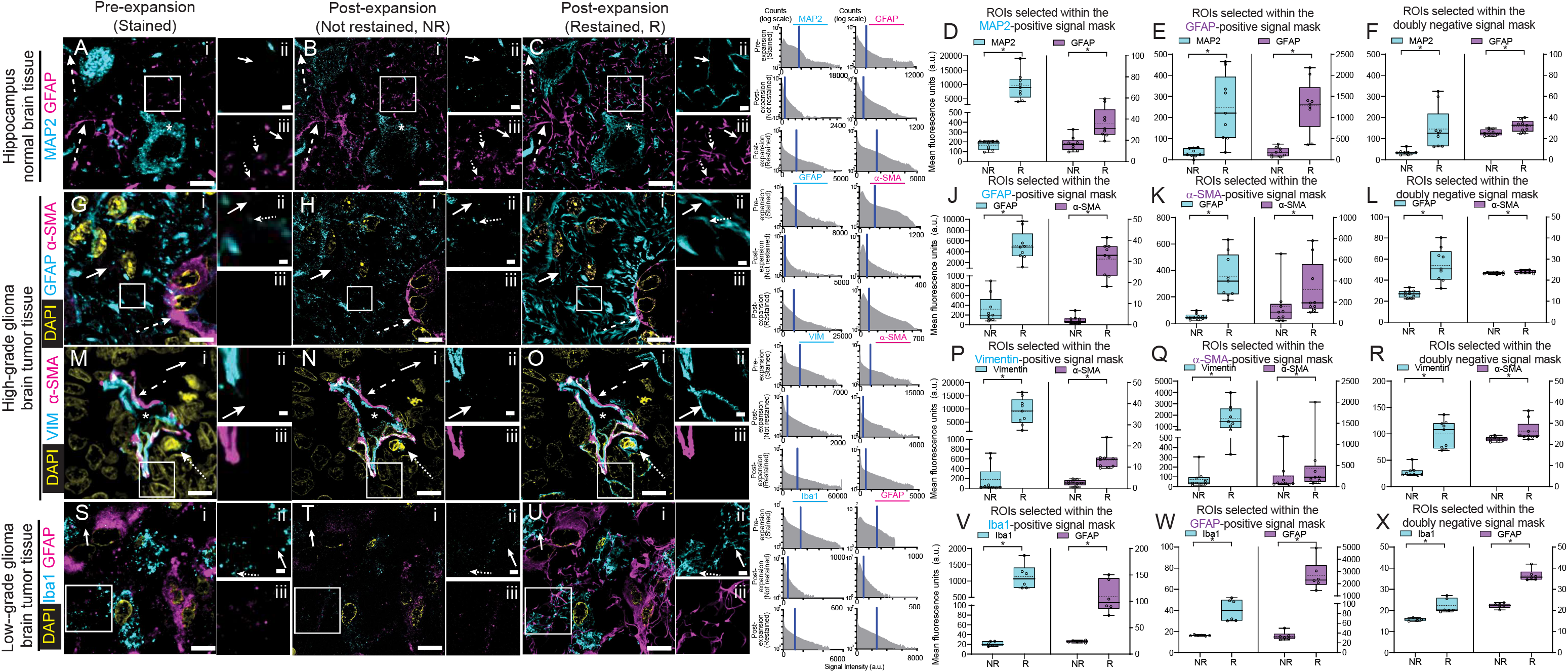
dExPath-enabled visualization of decrowded proteins reveals previously invisible cells and structures, compared to pre-expansion staining forms of expansion microscopy. (A) Representative (n = 3 samples, each from a different patient) pre-expansion confocal image (single z slice) of 5-μm-thick FFPE normal human hippocampal tissue. Sample underwent format conversion, antigen retrieval, and immunostaining for MAP2 and GFAP (processing as in Supp. Fig. 3A). White box in (i) marks a region with sparse and discontinuous signals that is shown magnified and in separate channels at the right (MAP2 in (ii) and GFAP in (iii)). MAP2 staining of a putative cell body (asterisk in (i)) and dendrite (upper dashed arrow in (i)). GFAP staining of a putative astrocytic process (lower dashed arrow in (i)) and discontinuous GFAP regions (dotted arrows in (iii)). Solid arrows show regions that were MAP2-negative (ii) or GFAP-negative (iii) in pre-expansion images (A), for comparison to postexpansion staining panels later in this figure. (B) Sample used for (A) after anchoring, gelation (Supp. Fig. 3B), softening (Supp. Fig. 3C), expansion (Supp. Fig. 3D) and imaging at ∼4x linear expansion. Sum intensity z-projection of an image stack covering the biological thickness of the original slice (used for all expanded images throughout this figure); images were of the same fields of view as in (A), using identical hardware settings. Asterisks and arrows as in (A). (C) Post-decrowding stained confocal images of the same fields of view as in (A-B) after decrowding and additional immunostaining for MAP2 and GFAP and re-expansion to ∼4x (Supp. Fig. 3E-F), again using identical hardware settings. Asterisks and arrows as in (A). For display purposes (A-C), histograms of pixel values for MAP2 and GFAP images were adjusted so that 1% of the pixels were saturated (histograms for A-C are shown in the subpanels, top to bottom). Vertical blue line, upper look-up table (LUT) limit (so that 1% of pixels are saturated). (D) For the entire set of images such as those of B and C, mean fluorescence intensities, from postexpansion with no additional staining (NR, “not restained”, as in B) and post-expansion restained (R, “restained”, as in C) images (raw image data, not adjusted as in the images of AC), averaged across MAP2-positive ROIs, for the MAP2 channel (cyan) and the GFAP channel (magenta). Box plot: individual values (open circles; 3 measurements were acquired from each patient), median (middle line), mean (dotted line), first and third quartiles (lower and upper box boundaries), lower and upper raw values (whiskers). Statistical testing: 2-tailed paired t-test (non-Bonferroni corrected) was applied to compare not restained vs. restained mean fluorescence intensities for each separate channel. *, p < 0.05. (E) As in D, but for GFAP-positive ROIs, for the MAP2 channel (cyan) and the GFAP channel (magenta). (F) As in D, but for doubly negative ROIs, for the MAP2 channel (cyan) and the GFAP channel (magenta). (G) Representative (n = 3 samples, each from a different patient) pre-expansion confocal image (single z slice) of 5-μm-thick FFPE human high-grade glioma tissue (cortex or white matter). Sample underwent format conversion, antigen retrieval, and immunostaining for GFAP and α-SMA (α-smooth muscle actin), and DAPI staining (Supp. Fig. 3A). White box in (i) marks a region with sparse and discontinuous signals that is shown magnified and in separate channels at the right (GFAP in (ii) and α-SMA in (iii)). α-SMA-staining of pericytes that are enveloping blood vessels (dashed arrow in (i)). Discontinuous GFAP regions (dotted arrow in (ii)). Solid arrows in (i) and (ii) show regions that were GFAP-negative pre-expansion (G), for comparisons to post-expansion staining panels later in this figure. (H) As in B, but for panel G. (I) As in C, but for panel G. (J) As in D, but for the GFAP (cyan) and α-SMA (magenta) channels, in GFAP-positive ROIs. (K) As in D, but for the GFAP (cyan) and α-SMA (magenta) channels in α-SMA-positive ROIs. (L) As in D, but for the GFAP (cyan) and α-SMA (magenta) channels in doubly negative ROIs. (M) Representative (n = 3 samples, each from a different patient) pre-expansion confocal image (single z slice) of 5-μm-thick human high grade glioma tissue (cortex or white matter). Sample underwent format conversion, antigen retrieval, and immunostaining for vimentin and α-SMA, and DAPI staining (Supp. Fig. 3A). White box in (i) marks a region including part of a blood vessel that is shown magnified and in separate channels to the right (vimentin in (ii) and α-SMA in (iii)). Vimentin and α-SMA-staining of the blood vessel wall (dashed arrow in (i)) which surrounds the vessel lumen (asterisk in (i)). A vimentin-positive cell outside the blood vessel (dotted arrow in (i)). Solid arrows in (i) and (ii) show regions that were vimentin-negative pre-expansion (M), for comparison to post-expansion staining panels later in this figure. (N) As in B, but for panel M. (O) As in C, but for panel M. (P) As in D, but for the vimentin channel (cyan) and the α-SMA channel (magenta), in vimentin-positive ROIs. (Q) As in D, but for the vimentin channel (cyan) and the α-SMA channel (magenta), in α-SMA-positive ROIs. (R) As in D, but for the vimentin channel (cyan) and the α-SMA channel (magenta), in doubly negative ROIs. (S) Representative (n = 3 samples, each from a different patient) preexpansion confocal image (single z slice) of 5-μm-thick human low grade glioma tissue (cortex or white matter). Sample underwent format conversion, antigen retrieval, and immunostaining for ionized calcium binding adapter molecule 1 (Iba1) and GFAP, and DAPI staining (Supp. Fig. 3A). White box in (i) marks a region with sparse and discontinuous signals that is shown magnified and in separate channels to the right (Iba1 in (ii) and GFAP in (iii)). Iba1 staining of discontinuous regions (dotted arrow in (ii)). Solid arrows in (i) and (ii) show regions that were Iba1-negative pre-expansion (S), for comparison to post-expansion staining panels later in this figure. (T) As in B, but for panel S. (U) As in C, but for panel S. (V) As in D, but for the Iba1 channel (cyan) and the GFAP channel (magenta), in the Iba1-positive ROIs. (W) As in D, but for the Iba1 channel (cyan) and the GFAP channel (magenta), in GFAP-positive ROIs. (X) As in D, but for the Iba1 channel (cyan) and the GFAP channel (magenta), in the doubly negative ROIs. Scale bars: (A-C) panel i, 9 μm; ii, 1.7 μm; (G-I) i, 7 μm; ii, 0.7 μm; (M-O) i, 8 μm; ii, 0.8 μm; (SU) i, 8 μm; ii, 0.8 μm. Linear expansion factors: (B,C) 4.1x; (H,I) 4.0x; (N,O) 4.3x; (T,U) 4.2x.

In one experiment (**Fig. 4A-C**), we used antibodies against the somato-dendritic marker, MAP2^35,59^ and the astrocytic marker, GFAP^36,57,58,60,61^. MAP2 staining yielded putative cell bodies and dendrites as well as sparser discontinuous dendrite-like regions (**Fig. 4A**). The latter regions remained as discontinuous puncta after 4x expansion (**Fig. 4B**). However, after post-expansion re-staining, new filaments appeared in areas that were previously completely MAP2-negative (**Fig. 4C**). Post-expansion staining not only improved the continuity of staining for existing structures, but reveals new, previously invisible, structures of appropriate morphology – as has been noted before in mouse brain tissue^24^, but now shown for human brain tissue. Similar improvements held for GFAP, with pre-expansion staining showing putative astrocytic processes as well as discontinuous signals (**Fig. 4A**). Post-expansion, resolution improved (**Fig. 4B**), and after re-staining, those regions appeared more continuous than pre-expansion (**Fig. 4C**). As with MAP2, completely new GFAP fibers also sometimes became visible, which were previously invisible.

To quantify the improvement in labeling post-expansion vs. pre-expansion, we constructed a binary image “signal” mask, for each stain, that corresponded to pixels that were positive (i.e., above a manually selected threshold) for a given stain in both pre-expansion as well as post-expansion staining images. (This method had the added bonus of excluding lipofuscin-positive pixels that would go dark in the post-expansion images, thus unnecessarily complicating interpretation). We also created a second “background” mask, for each stain, that corresponded to pixels that were negative (below the threshold mentioned before) in both the pre- and post-expansion staining images; a “doubly negative” background mask was also constructed, which corresponded to the pixels that were negative in both of the aforementioned background masks. Next, we constructed regions of interest (ROIs) that were small enough (0.2 microns) to fit entirely within the signal mask for a given stain, but that were at least an ROI width away from the signal mask for the other stain; we also constructed ROIs that were fully contained within the doubly negative mask, and similarly far from pixels that were positive in either signal mask. Finally, we calculated intensities averaged across the ROIs for the same locations in the expanded (**Fig. 4B**) vs. expanded-and-restained (**Fig. 4C**) images, to facilitate comparison. In regions positive in the MAP2 and GFAP signal masks (**Fig. 4D,** left and **Fig. 4E,** right), we saw increases of both signals in their respective ROIs (MAP2, p = 0.0003, 2-tailed paired t-test; n = 3 tissue samples from different patients; GFAP, p = 0.0007, 2-tailed paired t-test; n = 3 tissue samples from different patients), with mean intensity values going from ∼170 to ∼9400, and ∼190 to ∼1300 respectively. Of course, we would not expect MAP2 to occur much in GFAP-positive regions, nor GFAP in MAP2 regions. Thus these two genes give us the opportunity to assess whether post-expansion antibody application suffers from nonspecific staining. Indeed, GFAP, imaged under the same microscope settings that yielded the 190-1300 change in GFAP-positive regions, was ∼16 and ∼36 pre-expansion and post-expansion respectively (p = 0.0004, 2-tailed paired t-test; n = 3 tissue samples from different patients) in locations within the MAP2 signal mask, just a few percent of the biologically important (i.e., in the GFAP-positive ROIs) signals seen post-expansion. Similarly, MAP2, imaged in the GFAP signal mask, was ∼33 and ∼240 in pre- and post-expansion images (p = 0.003, 2-tailed paired t-test), again with intensities that are just a few percent of the biologically relevant signals seen post-expansion.

In the doubly negative regions, MAP2 intensities were ∼35 pre-decrowding and ∼150 post-decrowding, similar to the levels of MAP2 in the GFAP-positive area (**Fig. 4F,** left), and GFAP intensities were ∼25 pre-decrowding and ∼32 post-decrowding, similar to the GFAP levels in the MAP2-positive area (**Fig. 4F,** right). Thus, staining in the doubly negative regions is similar to that in the singly negative regions, further supporting the idea that the nonspecific staining is at levels that are a few percent of the biologically important signals, as noted in the previous paragraph.

We performed a similar analysis in high-grade glioma tissue from a human patient (**Fig. 4G-I**), staining for GFAP, which in glioma patients marks both astrocytes and glioma cells^57,62–64^, and α-SMA, a marker of pericytes^65–67^, which envelope blood vessels (**Fig. 4G**). As with MAP2 vs. GFAP, α-SMA and GFAP would not be expected to overlap, except perhaps at sites where astrocytes and glioma cells touch pericytes^67,68^; accordingly, we chose GFAP-positive and α-SMA-positive ROIs that were far apart from α-SMA and GFAP staining respectively, as well as doubly negative ROIs that exhibited neither. As before, GFAP became more continuous with post-expansion staining (**Figs. 4G-I**), showing new filaments, and an overall increase in intensity in GFAP-positive ROIs (**Fig. 4J**, left; pre, ∼320 vs post, ∼5100; p = 0.0006, 2-tailed paired t-test; n = 3 tissue samples from different patients). α SMA intensity also went up in α SMA-positive regions (**Fig. 4K**, right; pre, ∼160 vs post, ∼320, p = 0.0006 2-tailed paired t-test; n = 3 tissue samples from different patients). In contrast, α SMA was ∼2 and ∼30, pre- and post-expansion respectively, in GFAP-positive ROIs (**Fig. 4J**, right; p = 0.004, 2-tailed paired t-test; n = 3 tissue samples from different patients); GFAP was ∼50 and ∼350, pre- and post-expansion, in α MA - positive ROIs (p = 0.004; 2-tailed paired t-test; n = 3 tissue samples from different patients) – in each case, a small fraction of the biologically important signals measured in the appropriate ROI. And GFAP and α-SMA values in the doubly negative ROIs were comparably low (**Fig. 4L**).

As a third check, we examined vimentin and SMA in high-grade glioma tissue. Vimentin is expressed in some tumor cells^69^, some activated microglia^70^, as well as all endothelial cells^37^ and some pericytes^71^. Thus, vimentin would be expected to sometimes be near, or even -SMA (i.e., in pericytes) and sometimes to be well-isolated from α in other cell types) ^67,68,72^. We observed vimentin and α-SMA signals in the blood vessel wall and surrounding the vessel lumen (**Fig. 4M**). Vimentin signals were also observed in cells (e.g., putative tumor cells or activated microglia) outside of blood vessels (**Fig. 4M**); with similar observations after 4x expansion (**Fig. 4N**). However, after post-expansion re-staining (**Fig. 4O**), new vimentin-positivity appeared in cells, far from blood vessels, that were previously vimentin-negative (**Fig. 4M-O**). We analyzed vimentin ROIs far away from α SMA, and found the vimentin staining to go up from ∼170 to ∼9100 in these ROIs (**Fig. 4P**, left; p = 0.0008, 2-tailed paired t-test; n = 3 tissue samples from different patients); in α SMA ROIs, we found vimentin was found in these ROIs, and vimentin also went up significantly, from ∼80 to ∼1750 (**Fig. 4Q**, left; p = 0.0001, 2-tailed paired t-test; n = 3 tissue samples from different patients), as expected. In contrast, α SMA was very little located in the vimentin ROIs (∼3 and ∼13, pre- and post-expansion; **Fig. 4P**, right; p = 0.0001, 2-tailed paired t-test; n = 3 tissue samples from different patients), and went up in α SMA ROIs (from ∼240 to ∼440; **Fig. 4Q**, right; p = 0.04, 2-tailed paired t-test; n = 3 tissue samples from different patients) to some extent – clearly, not all proteins are equally crowded in all cells; perhaps α SMA is relatively uncrowded to begin with. As before, doubly negative staining was consistently low (**Fig. 4R**).

Finally, as a fourth check, we examined ionized calcium binding adapter molecule 1 (Iba1) and GFAP in low-grade glioma tissue, again from cortex or white matter. Iba1 is expressed in macrophages and microglia^73^. The places we would expect colocalization of these two markers are at sites where an Iba1-positive cell (i.e., macrophage, microglia)^73^ and a GFAP-positive cell (i.e., astrocyte, glioma) touch^74^, or where microglia have phagocytosed GFAP-containing fragments^74^, or possibly a cell type with a dual astrocytic and macrophage/microglia molecular phenotype^75–78^. Accordingly, we chose ROIs that were Iba1-positive or GFAP-positive that were far apart from GFAP and Iba1 staining respectively, as well as doubly negative ROIs that exhibited neither. We observed GFAP and Iba1 signals in distinct cells before (**Fig. 4S**) and after 4x expansion (**Fig. 4T**). However, after post-expansion re-staining (**Fig. 4U**), new Iba1-positivity appeared in regions that were previously Iba1-negative (**Fig. 4S-U**), and generally appeared more continuous (**Fig. 4S-U**). Iba1 increased in intensity in Iba1-positive ROIs (**Fig. 4V**, left; pre, ∼21 vs post, ∼1100; p = 0.0009, 2-tailed paired t-test; n = 3 tissue samples from different patients). GFAP also went up in GFAP-positive regions (**Fig. 4W**, right; pre, ∼34 vs post, ∼2700, p = 0.003 2-tailed paired t-test; n = 3 tissue samples from different patients). In contrast, GFAP was ∼24 and ∼110, pre- and post-expansion respectively, in Iba1-positive ROIs (**Fig. 4V**, right; p = 0.0009, 2-tailed paired t-test; n = 3 tissue samples from different patients); Iba1 was ∼16 and ∼40, pre- and post-expansion, in GFAP-positive ROIs (**Fig. 4W**, left; p = 0.002; 2-tailed paired t-test; n = 3 tissue samples from different patients) – in each case, a small fraction of the biologically important signals measured in the appropriate ROI. And as before, the Iba1 and GFAP values in the doubly negative ROIs were comparably low (**Fig. 4X**).

Having validated the decrowding aspect of dExPath, we next examined whether the improved immunostaining facilitated by dExPath improved images vs. those obtained by an earlier, similar-resolution super-resolution method that does not decrowd epitopes, SR-SIM. We first performed antigen retrieval and stained high-grade glioma and normal hippocampus with anti-vimentin or anti-MAP/anti-GFAP, using in both cases DAPI to counterstain the cells’ nuclei. Samples were imaged by SR-SIM (**Fig. 5A,B**), followed by the first part of the dExPath protocol (i.e., chemical softening and expansion) (**Supp Fig. 3A-D**) to acquire confocal images post-expansion but with pre-decrowding-staining (**Fig. 5C-D**). Next, we performed the last part of the dExPath protocol (i.e., post-decrowding staining) (**Supp Fig. 3E-F**) to acquire confocal images post-expansion with post-decrowding-staining (**Fig. 5E-F**). Both SR-SIM and post-expansion confocal images of pre-decrowding-stained tissue revealed highly punctate patterns for vimentin (**Fig. 5A, 5C**) as well as MAP2 and GFAP (**Fig. 5B,5D**). In contrast, all these stains revealed continuous structures in confocal images taken after post-decrowding staining (**Fig. 5E, 5F**), as well as new filamentous structures that had not been previously observed (compare **Fig. 5A,C,E** vs. **5B,D,F**). Thus, the improvement in staining continuity and revelation of new structures that was borne out by our pre- vs. post comparisons are also manifest as improvements over pre-expansion staining super-resolution. dExPath may provide a general solution to the problem of punctate staining appearances in brain tissues, when they should appear continuous, in super-resolution microscopy^2,9,11,13,79^.

**Fig. 5.**
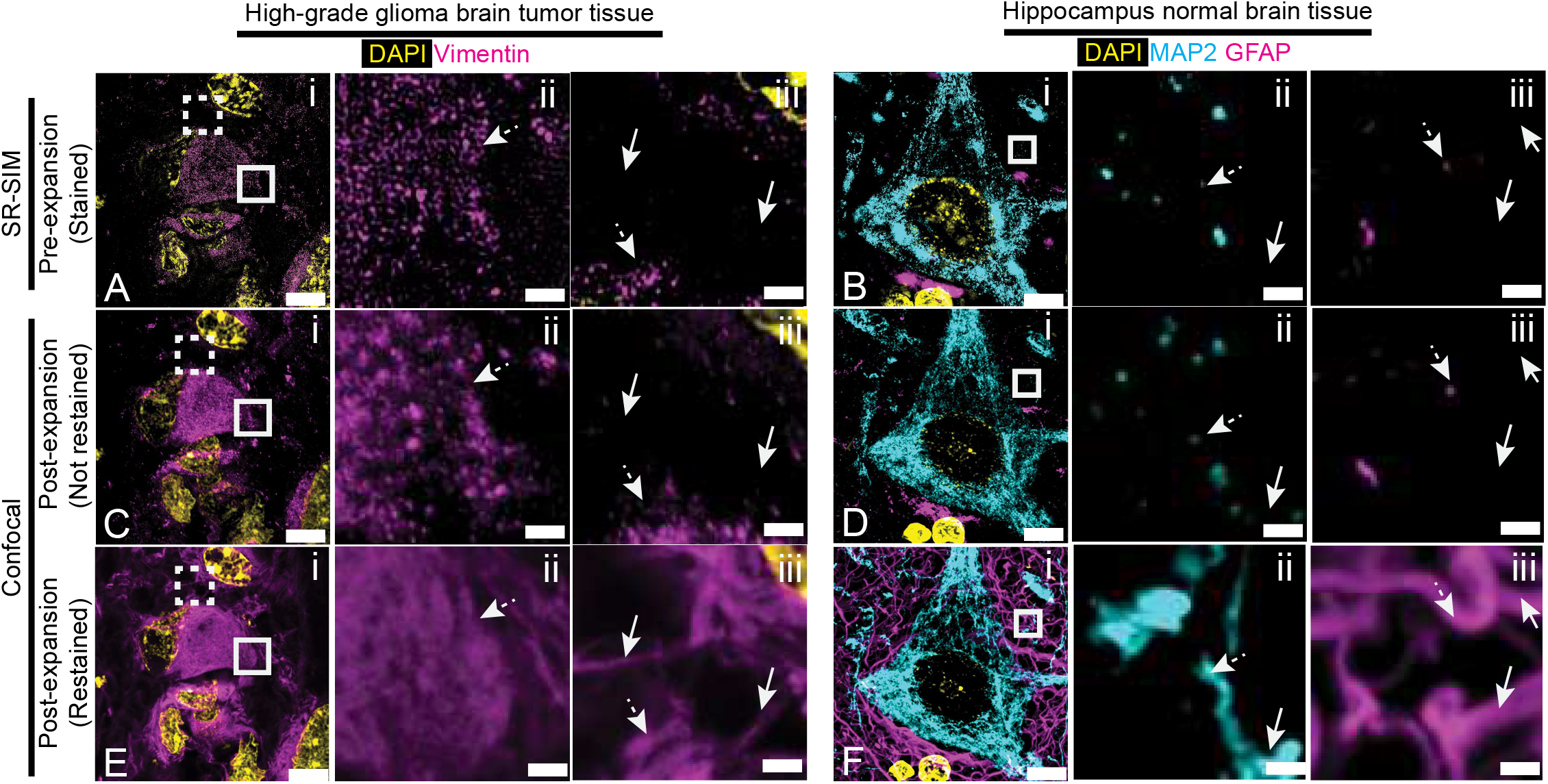
dExPath-enabled visualization of decrowded proteins reveals previously invisible cells and structures, compared to SR-SIM imaging of unexpanded tissues. (A-B) Representative pre-expansion SR-SIM images of FFPE 5-µm-thick human high-grade glioma tissue (A, n = 3 samples, each from a different patient) and normal human hippocampus (B, n = 3 samples, each from a different patient). Samples underwent format conversion, antigen retrieval, and immunostaining for vimentin, and DAPI staining (A), or immunostaining for MAP2 and GFAP, and DAPI staining (B) (processing as in Supp. Fig. 3A). (A) Solid and dashed white boxes in (i) mark two separate regions shown magnified in (ii) (solid box) and (iii) (dashed box), respectively. Dotted arrows mark regions that appear as punctate and discontinuous in pre-expansion SR-SIM images for vimentin in (ii) and (iii), and solid arrows mark regions that were negative for vimentin in (iii), for comparison to post-expansion staining panels later in this figure. (B) Solid white box in (i) shown magnified in (ii) for MAP2 and in (iii) for GFAP. Arrows as in (A) but for MAP2 and GFAP, in their respective images. (C-D) Samples used for (A-B) after anchoring, gelation (Supp. Fig. 3B), softening (Supp. Fig. 3C), expansion (Supp. Fig. 3D) and confocal imaging at ∼4x linear expansion. Sum intensity z-projection of an image stack covering the biological thickness of the original slice (used for all expanded images throughout this figure); images were of the same fields of view as in (A-B). Arrows as in (A-B). (E-F) Post-decrowding stained confocal images of the same fields of view as in (A-B) after decrowding and additional immunostaining for vimentin (E), or MAP2 and GFAP (F), followed by DAPI staining and re-expansion to ∼4x (Supp. Fig. 3E-F), imaged using identical hardware settings as in (C-D). Arrows as in (A-B). Brightness and contrast settings in images (A-F): first set by the ImageJ auto-scaling function, and then manually adjusted (by raising the minimum-intensity threshold and lowering the maximum-intensity threshold) to improve contrast for stained structures. Scale bars (in biological units): (A, C, E) left column, 8.3 µm; middle and right columns 840 nm; (B, D, F) left column, 6.0 µm; middle and right columns 500 nm. Linear expansion factors: (C) 4.1x; (D) 4.3x; (E) 4.1x; (F) 4.2x

### dExPath-mediated visualization of protein targets in the mouse brain tissue

dExPath worked well on mouse brain tissue, fixed through standard PFA fixation protocols, and resulted in high quality images of mouse cortex and white matter, using antibodies against proteins including bassoon, synaptophysin, homer, post-synaptic density protein 95 (PSD95), MAP2, histone, MBP^36,80^, neurofilament medium chain (NF-M), SMI-312, neurofilament light chain (NF-L)), GFAP, giantin, laminin, collagen IV^81^, and zona occludens-1 (ZO-1)^82^. Pre- and post-synaptic markers often were localized near each other (**Fig. 6A, 6B**). MBP staining revealed tubular structures, appropriate for putative axons (**Fig. 6C**). Staining intermediate filament proteins (NF-M, NF-L, SMI-312 and GFAP) resulted in expected patterns (**Fig. 6D, E**). Staining for components of blood vessels, laminin, collagen IV, and ZO-1, showed patterns reminiscent of blood vessels (**Fig. 6F**). Thus, we show that dExPath provides an avenue for generally decrowded nanomapping of proteins in PFA-fixed brain tissue. In addition, we also demonstrated the ability of dExPath to enable multiple rounds of staining on the same tissue samples, allowing for a high degree of decrowded protein multiplexing in human tissues at nanoscale resolution (**Supp. Figs. 7** and **8**).

**Fig. 6.**
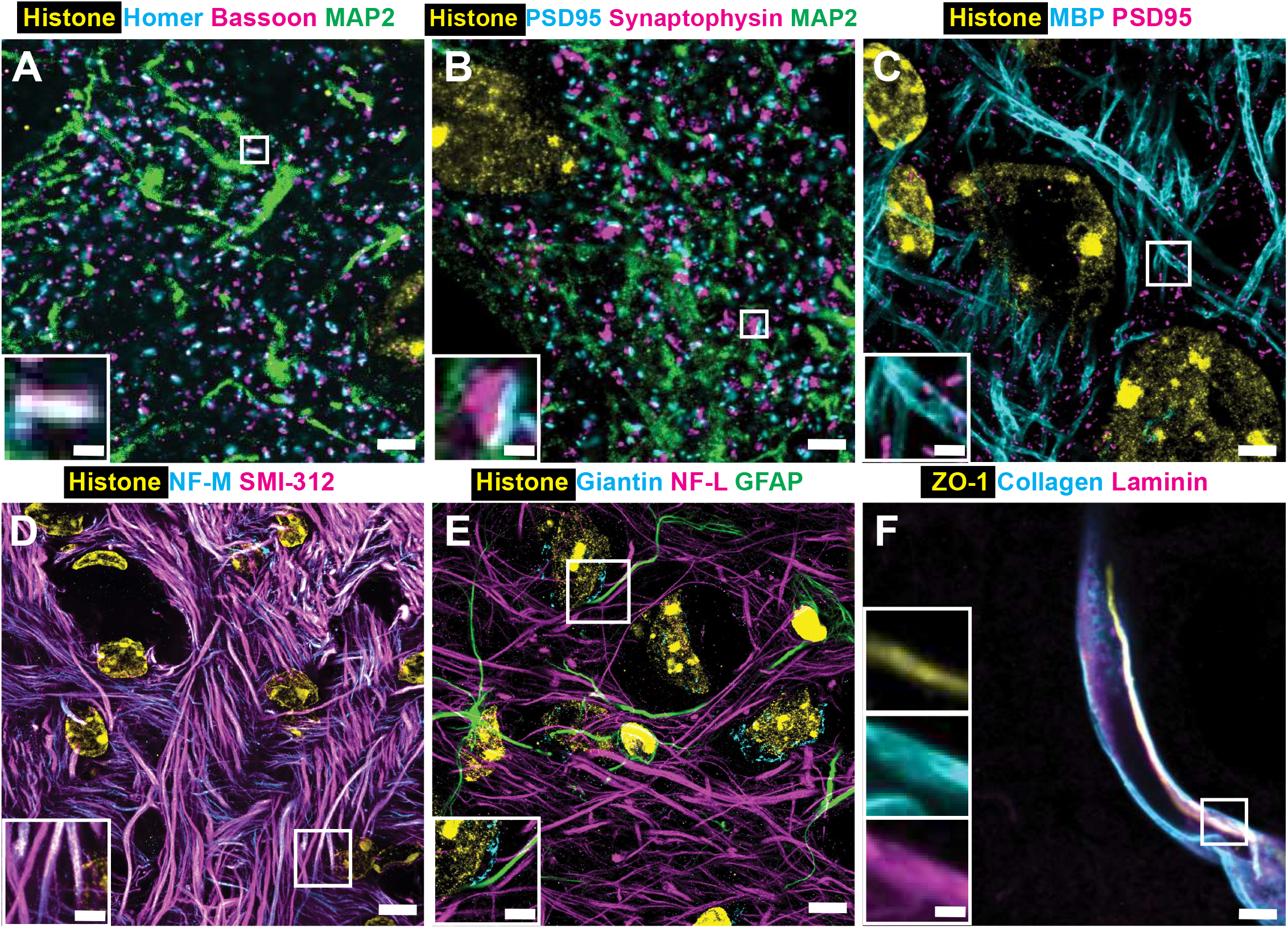
dExPath applied to formaldehyde-fixed mouse brain tissue. (A-F) Confocal images of 4%-PFA-fixed, 10-µm-thick samples of mouse cortex. Samples underwent format conversion (Fig. 1A), anchoring, gelation (Fig. 1B), softening (Fig. 1C), de-crowding (Fig. 1D), post-decrowding immunostaining, and confocal imaging at 4x linear expansion (Fig. 1E). The tissue samples were stained for the following: (A) histone H3 (a nuclear protein), homer (a postsynaptic protein), bassoon (a presynaptic protein), and MAP2. (B) histone H3, postsynaptic density protein 95 (PSD95, a postsynaptic protein), synaptophysin (a presynaptic protein), and MAP2. (C) histone H3, myelin-basic protein (MBP, a protein of myelinated axons), and PSD95. (D) histone H3, neurofilament medium chain (NF-M, a neurofilament subunit), and SMI-312 (a pan-axonal marker of neurofilaments). (E) histone H3, giantin (a protein of the Golgi complex), neurofilament light chain (NF-L, a neurofilament subunit distinct from NF-M), and GFAP. (F) zona occludens-1 (ZO-1, a protein of tight junctions), and laminin and collagen IV (two distinct proteins of the basement membrane of blood vessels). White boxes mark regions shown magnified in insets on the left. All images are sum intensity z-projections of a confocal image stack. Brightness and contrast settings in images (A-F): first set by the ImageJ auto-scaling function, and then manually adjusted (by raising the minimum-intensity threshold and lowering the maximum-intensity threshold) to improve contrast of stained structures. Scale bars (in biological units): (A-B) outer panel, 2.25 µm; inset, 300 nm; (C) outer panel, 2.5 µm; inset, 500 nm; (D) outer panel, 6 µm; inset, 1.25 µm; (E) outer panel, 6 µm; inset, 2 µm; (F) outer panel, 1.45 µm; inset, 400 nm; Linear expansion factors: (A) 4.0 x; (B) 4.1 x; (C) 4.2 x; (D) 4.0 x; (E) 3.9 x; (F) 4.2 x.

### dExPath reveals cell populations exhibiting combinations of disease-state markers in human glioma tissue

Our prior experiment using glioma tissues (**Figs. 4** and **5)** demonstrated that post-expansion staining increases the intensity, continuity, and number of structures stained for vimentin, Iba1 and GFAP compared to pre-expansion staining. Therefore, we next asked whether this could lead to detecting more cells carrying specific antigen combinations, which might alter interpretation of clinical biopsies as well as basic understanding of brain tumor biology. For example, a cell exhibiting GFAP alone would be considered an astrocyte or a tumor cell, but a cell with both GFAP and vimentin would be considered a tumor cell with more aggressive features than a vimentin-negative/GFAP-positive tumor cell^83–86^. If, with post-expansion staining, we saw more dually labeled cells than before, then more cells than previously thought may be aggressive tumor cells, or perhaps the aggressive tumor cells traditionally studied are a subpopulation of the entire set of such cells, with the newly discovered cells perhaps representing a subpopulation of aggressive tumor cells with different properties^86,87^.

We imaged low-grade glioma tissue sections serially 1) after antigen retrieval and pre-expansion immunostaining (**Fig. 7A**); 2) after dExPath softening, washing with PBS (which results in an expansion factor of ∼2.3x), tissue shrinkage (via adding salt to expansion factor of 1.3x, **Fig. 7B**); 3) after ∼4x expansion (∼4x, **Fig. 7C**); 4) after post-decrowding immunostaining, washing (∼2.3x) and shrinkage (∼1.3x) (**Fig. 7D**); and 5) after a final expansion step back to ∼4x (**Fig. 7E**). We then compared the initial pre-expansion immunostained state (**Fig. 7A**) to the post-decrowded immunostained state at ∼1.3x (**Fig. 7D**), because in this analysis we are concerned primarily with determining if two stains are found in the same cell, as opposed to the nanoscale emphasis in **Fig. 4** of mapping fine-scale, e.g. filamentous, structures; in addition, we were curious to know if, independently of the improved resolution of expansion microscopy, the enhanced staining afforded by decrowding could reveal new clinically relevant features, even without significant physical magnification.

**Fig. 7.**
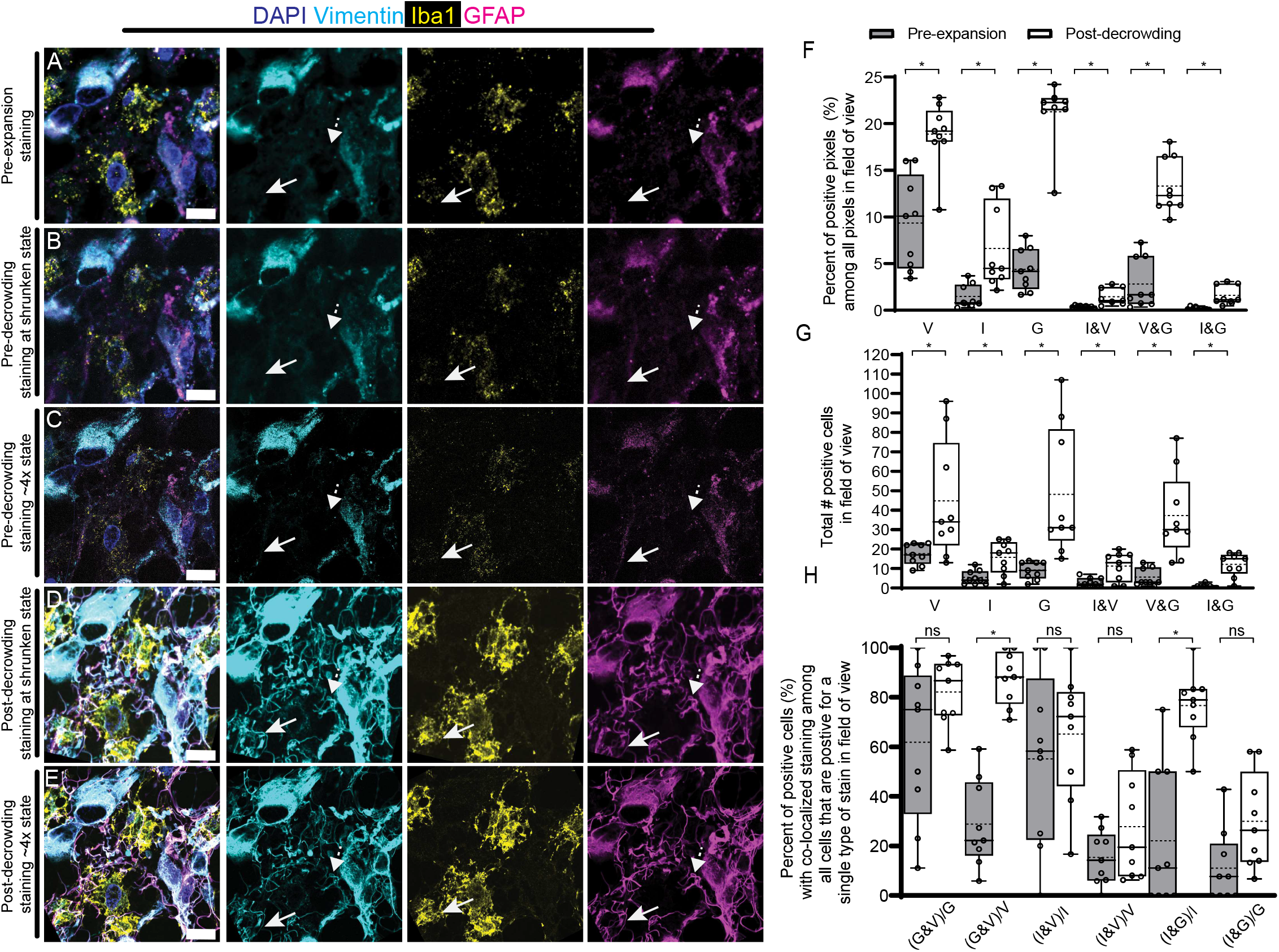
dExPath reveals large numbers of previously undetected cells defined by single and multiple markers of importance to glioma biology. (A) Representative (n = 3 samples, each from a different patient) pre-expansion confocal image (single z slice) of a 5-µm-thick FFPE human low-grade glioma specimen. Sample underwent format conversion, antigen retrieval, immunostaining for vimentin, Iba1 and GFAP, and DAPI staining (Supp. Fig. 3A). Left panel, overlay of all 4 channels; right three panels, individual channels (not including DAPI). Dotted arrows show regions that were vimentin and GFAP negative in pre-expansion images, and solid arrows show regions that were Iba1, GFAP and vimentin negative in pre-expansion images, for comparison to post-decrowding staining panels later in this figure. (B) Sample used for (A) after anchoring, gelation (Supp Fig. 3B), softening (Supp Fig. 3C), washing with PBS (which results in an expansion factor of ∼2.3x), tissue shrinkage (via adding salt) to ∼1.3x of the original size, and confocal imaging. Single z slice image centered at the same midpoint of the original slice; images were of the same field of view as in (A), using identical hardware and software settings. Arrows as in (A). (C) Sample used for (B) after expansion (Supp. Fig. 3D) and confocal imaging at ∼4x linear expansion. Sum intensity z-projection of an image stack covering the biological thickness of the original slice; images were of the same field of view as in (A), using identical hardware and software settings. Arrows as in (A). (D) Post-decrowding stained confocal images of the same field of view as in (A) after decrowding and additional immunostaining for vimentin, Iba1, and GFAP, tissue shrinkage (Supp. Fig. 3E-F) and confocal imaging at shrunken state. Arrows as in (A). (E) Sample used for (D) after expansion (Supp. Fig. 3D) and confocal imaging at ∼4x linear expansion. Arrows as in (A). (F) For the entire set of images such as those for (A) and (D), pixel level analysis of the percent of singly or doubly positive stained pixels, from pre-expansion (gray boxes) and post-decrowding at shrunken state (white boxes) images, for vimentin (V), Iba1 (I), GFAP (G), Iba1 and vimentin (I&V), vimentin and GFAP (V&G), and Iba1 and GFAP (I&G). Values represent percent of positive pixels among all pixels in the field of view. with 3 different values per sample each corresponding to a different field of view Box plot: individual values (open circles; 3 measurements were acquired from each patient), median (middle line), mean (dotted line), first and third quartiles (lower and upper box boundaries), lower and upper raw values (whiskers), used throughout the graphs of this figure. (G) As in (F), but with cell level analysis of the total number of singly or doubly positive labeled cells, from pre-expansion and post-decrowding at shrunken state images. Values represent total number of labeled cells in the field of view. (H) As in (G) but with cell level analysis of the percentage of doubly positive labeled cells (two stains separated by “&” in the x-axis) divided by all singly positive cells for a stain (single stain shown after the slash in the x-axis) in the pre-expansion and post-decrowding at shrunken state images. Values represent the percentage (%) of doubly positive cells relative to the total number of singly positive cells for a stain. Brightness and contrast settings in images (A-E): first set by the ImageJ auto-scaling function, and then manually adjusted (by raising the minimum-intensity threshold and lowering the maximum-intensity threshold) to improve contrast for stained structures but quantitative analysis in (F-H) was conducted on raw image data. Statistical testing: 2-tailed paired t-test (non-Bonferroni corrected) were applied on pre-expansion and post-decrowding values. *, p < 0.05; ns, not significant. Scale bars: (A-E) 11 µm. Linear expansion factors: (B, D) 1.3x; (C, E) 4.4x.

By comparing samples in the pre-expansion and shrunken ∼1.3x state (**Fig. 7A, 7B, 7D**), we observed that post-decrowding immunostaining (**Fig. 7D**) was able to reveal additional vimentin-, GFAP-, and Iba1-positive staining not detected in the pre-expansion (**Fig. 7A**) or pre-decrowding (**Fig. 7B**) states, as expected from the improvements shown in **Fig. 4.** In addition, some regions showed increased signal, after post-decrowding immunostaining, for multiple molecular species. For example, some regions showed new structures that were GFAP- and vimentin-positive (compare **Fig. 7D–7E** vs. **7A–7C**), or Iba1, GFAP and vimentin positive (compare **Fig. 7D–7E** vs. **7A–7C**), which were not visible with pre-expansion staining. Indeed, when we examined the fraction of pixels that were positive for each individual stain in single z-slices of pre-expansion (**Fig. 7A**) and post-decrowding (**Fig. 7D**) images, they increased significantly, and furthermore, the fraction of pixels positive for multiple molecular species increased as well (**Fig. 7F**).

These increases in stain-positive pixels translated into significant increases in the number of individual cells identified with either single or multiple labels (**Fig. 7G**). The number of cells positive for vimentin went up almost 3-fold, the number of cells positive for GFAP went up over 5-fold, and the number of cells positive for Iba1 went up almost 3-fold (vimentin, p = 0.032, 2-tailed paired t-test, n = 3 tissue samples from different patients; GFAP, p = 0.0071, 2-tailed paired t-test, n = 3 tissue samples from different patients; Iba1, p = 0.0011, 2-tailed paired t-test, n = 3 tissue samples from different patients). Thus, the number of cells corresponding to some tumor cells, some activated microglia, as well as all endothelial cells and some pericytes of mesenchymal origin (vimentin), or astrocytes and glioma cells (GFAP), or macrophages and microglia (Iba1), increased dramatically, suggesting that many cell types important for glioma pathology and response may be quantitatively underestimated by conventional immunostaining.

As mentioned earlier, a cell with both GFAP and vimentin is an aggressive tumor cell cell^83–86^, a cell with Iba1 and vimentin is an activated macrophage or microglial cell^70,73,88^, and a cell with Iba1 and GFAP is either a macrophage or microglial cell that phagocytosed a GFAP expressing cell (astrocyte or tumor cell) or a cell type with a dual astrocytic and macrophage/microglia molecular phenotype^74–76,78,82^. In each case, the dually labeled cell is qualitatively different from a singly labeled one. Cells positive both for GFAP and vimentin, identified as aggressive/invasive tumor cells, increased in number by about 6-fold with post-expansion vs. pre-expansion staining, suggesting that many more aggressive/invasive tumor cells are present than previously thought (**Fig. 7G,** p = 0.0035, 2-tailed paired t-test, n = 3 tissue samples from different patients). Amongst GFAP-expressing cells, we observed a ∼30% increase in the fraction that were vimentin-positive (**Fig. 7H**, p = 0.036, 2-tailed paired t-test, n = 3 tissue samples from different patients), suggesting that even in low-grade gliomas, a vast majority of tumor cells may be aggressive. Cells double-labeled with Iba1 and vimentin increased by about 4-fold using post-expansion vs. pre-expansion staining (**Fig. 7G,** p = 0.0030, 2-tailed paired t-test, n = 3 tissue samples from different patients), suggesting that a majority of activated macrophages and microglia might currently be overlooked.

Similarly, cells double-labeled with Iba1 and GFAP increased by about 10-fold with post-expansion vs pre-expansion staining (**Fig. 7G,** p = 0.00043, 2-tailed paired t-test, n = 3 tissue samples from different patients). These dual labeled cells are indicative of two cell populations. One population is that of macrophages or microglia, which have phagocytosed GFAP-expressing cells or debris in the tumor tissue sample (e.g., from astrocytes or tumor cells)^74–76,78,82^. Macrophage or microglial phagocytosis of GFAP-expressing cells and their debris may support tumor growth via removing of debris such as apoptotic corpses from the tumor microenvironment^74^. The second cell population might be a cell population found in diseased states such as stroke and neurodegenerative states^75,76,82^ and recently found to be present in glioblastoma^78^, in which cells share the molecular signatures of both Iba1 expressing cells (macrophages or microglia) and GFAP expressing cells (astrocytes or tumor cells) ^74,78^. We show a substantial increase of these Iba1-GFAP dually labeled cells, which can have a protumorigenic role in low-grade gliomas^74,78^. Approximately 80% of Iba1-expressing cells also exhibited GFAP post-expansion, versus only 20% pre-expansion (p = 0.000094, 2-tailed paired t-test, n = 3 tissue samples from different patients, **Fig. 7H**). In summary, we observed a significant increase in the percentage of immune cells with phenotypes of importance for the growth of low-grade gliomas.

Taken together, our post-expansion staining revealed “undercover” aggressive features in tumors traditionally considered indolent or mildly aggressive. These results suggest that post-expansion staining could uncover cell populations that were either present but not accounted for, or new cell populations with more aggressive features than previously thought, thus increasing clinico-pathological accuracy.

## Discussion

We describe here a new form of expansion microscopy, dExPath, that enables immunostaining of decrowded proteins, for nanoscale level visualization of previously unseen biological structures and cell populations in human clinical tissue specimens. The innovation of our method rests on the discovery that isotropic magnification of tissues, together with antigen preservation, enables protein decrowding in human tissues, addressing a fundamental problem of immunostaining: the inaccessibility of target epitopes by antibodies due to their physical size^3–5,9–12,89^. We found that dExPath works across both normal and diseased brain tissue (e.g., low- and high-grade gliomas) types, and improves immunostaining for many molecular targets. We showed that dExPath enables immunostaining of previously inaccessible cells or subcellular features in normal brain and tumor tissues, showing the potential for dExPath as a tool for clinicians and researchers to uncover immunostaining patterns previously unseen in diseased tissues for improved diagnostics and analysis of tissue architecture.

A potential mechanistic explanation for these improvements is that post-decrowding staining increases the number of spatially accessible epitopes on the target protein, which results in an increased labeling density of the antibodies and their associated fluorescent signal. This hypothesis aligns well with previous studies that demonstrate improvement in immunostaining by using small-sized probes (e.g., ∼3nm) due to improved probe access to targets^3,4,9,13,14,79^. While the type of improvements could be similar between dExPath and the small-sized-probe approach, dExPath directly supports the vast library of conventional off-the-shelf antibodies, which circumvents the need for synthesizing specialized probes, and can therefore be directly applied immediately in academic and clinical settings.

We discovered that dExPath has the unexpected capability to remove the intense autofluorescence from lipofuscin aggregates found in senescent brain tissues^42–49^, improving the accuracy of IHC-mediated detection of intracellular structures. While other methods exist for the masking or quenching of lipofuscin autofluorescence, such as with Sudan Black B^50^, they have also been associated with limitations including interruption of antibody binding, and reduction of on-target fluorescence^43,45,48,49^, which are in stark contrast to the enhanced immunostaining results from dExPath.

In addition dExPath provides a robust method for highly multiplexed immunostaining of decrowded proteins, by retaining protein antigenicity and high isotropy of tissue microarchitecture across sequential rounds of antibody stripping and re-staining. These capabilities could be particularly useful for mapping complex cellular and molecular types in both normal and diseased tissue microenvironments.

This study examined several antigen targets that have been commonly used as molecular markers to identify specific cell types or cell states that are important in normal or diseased brains, which ties the additional features that we observed from dExPath-mediated post-expansion immunostaining to potential clinical significance, deserving of further investigation beyond the scope of this technology-focused paper. For example, we showed that dExPath not only revealed abundant GFAP-positive filaments in non-diseased human brain tissue, via its decrowding capability, but these filaments can also be clearly resolved, via its capability to perform facile volumetric nanoscale imaging (∼70 nm, at 4x linear expansion factor). GFAP is involved in multiple physiological and injury-induced functions in which the precise mechanism of this protein remains unknown but its spatial localization appears critical for function (for example, formation of glial scars^90 91^, maintenance of myelinated sites^92^, lining of the blood-brain barrier ^93^, etc.). Accordingly, our improved capability to visualize the network of GFAP-positive filaments would likely facilitate studies of neurobiology and cellular responses to brain injury in the highly clinically relevant human context.

Our triple staining experiment (vimentin, Iba1, GFAP) of low-grade glioma tissues showed that dExPath can reveal substantially increased colocalization between these cell type markers, with great implications for the analysis of different cell populations in glioma biology. For example, our detection of a significant number of previously undetected double-labeled GFAP- and vimentin-positive cells in low-grade glioma tissue may represent a nascent indication of a malignant cell subpopulation in these tumors^94–98^, usually not detected histologically. Similarly, cells double-labeled with Iba1 and vimentin (interpreted as activated macrophages or microglial cells^70,73,88^) may represent a smoldering status of immune activation that could have major clinical relevance in these tumors, and cells double-labeled with Iba1 and GFAP may represent a large increase in the number of phagocytic macrophages/microglia, or possibly an increase in tumor cells with phagocytic properties with an increased invasive ability^73,77,78 99–101^.

The significant cellular details provided by dExPath, coupled with its compatibility with archival and routine pathology samples, make this technique particularly powerful to aid in improved analysis of malignant brain tumors, with the potential, down the line, to serve in diagnostic capabilities once sufficient clinical familiarity with dExPath has occurred. While we have primarily focused on glioma tissues for this study, dExPath could be applied to other malignancies or neuropathologies, helping in the short term with detailed scientific analysis of such conditions, and in the future, therapeutic decision-making by more thoroughly revealing disease specific molecular markers and by facilitating multiplexed readout of these markers from minimal amounts of biopsied tissue.

dExPath achieves protein decrowding and highly multiplexed immunostaining of clinical samples while enabling nanoscale resolution imaging on conventional microscopes, all accomplished using low cost, commercially available reagents and instruments found in a conventional basic science or pathology laboratories. We anticipate broad utility of dExPath in many scientific and clinical contexts.

## Methods

### Human and animal samples

The normal brain, low- and high-grade glioma human samples used in this study were all 5-µm-thick formalin-fixed paraffin-embedded (FFPE) tissue microarrays bought from US Biomax. The use of unused, unidentified archival specimens does not require informed consent from the subjects.

All procedures involving animals were conducted in accordance with the US National Institutes of Health Guide for the Care and Use of Laboratory Animals and approved by the Massachusetts Institute of Technology Committee on Animal Care. Male and female 12-16 weeks old, wild type (Swiss Webster) mice were used in this study. Mice were deeply anesthetized with isoflurane and perfused transcardially with ice cold phosphate buffered saline (PBS) followed by ice cold 4% paraformaldehyde (PFA) in phosphate buffered saline (PBS). Brains were harvested and postfixed in the same fixative solution at 4°C overnight. Fixed brains were incubated in 100 mM glycine for 1-2hrs at 4°C and sectioned to 10 um-thick slices with a vibratome (Leica VT1000S).

### Tissues processing methods

#### Format conversation, antigen retrieval, and pre-expansion immunostaining

For FFPE 5-µm thick samples of normal human hippocampus or cortex and human low- or high-grade glioma brain tumor tissues, format conversion (**Fig. 1A; Supp. Fig. 1A; 2A; 3A; 4A**) entails deparaffinization and rehydration, which includes 2 washes in 100% xylene for 3 min each, and then serial incubation in the following solutions, for 3 min each and all at room temperature (RT): (1) 50% xylene + 50% ethanol, (2) 100% ethanol, (3) 95% ethanol (in deionized water, as for all the following ethanol dilution solutions), (4) 80% ethanol, (5) 50% ethanol, (6) deionized water, and (7) 1x PBS (**Fig. 2A,B; 3A-C,L; 4A,G,M,S; 5A,B; 7A**). For 4%-PFA 10-µm thick samples of normal mouse brains, format conversion entails 3 washes in 1x PBS at RT for 5 min each (**Fig. 6**).

Following format conversion, tissues samples were designated for 1) pre-expansion immunostaining (**Supp. Fig. 3A** for **Fig. 2A,B; 3L; 4A,G,M,S; 5A,B; 7A;** and **Supp. Fig. 4A** for **Supp. Fig. 5A,B**); 2) no pre-expansion immunostaining and only pre-expansion DAPI staining at 2 µg/ml in 1 x PBS at RT for 15 min (**Fig. 1A**) (**Fig. 3A-C; Supp. Fig. 6A,B**); 3) or directly to the next steps in our protocol (**Fig. 1B-E; Supp. Fig. 2B-F**) without any pre-expansion staining (**Fig. 6; Supp Fig. 7; 8**).

For tissue samples that were designated for pre-expansion immunostaining, following format conversion, we applied antigen retrieval to enhance immunostaining^6,40,41^. Antigen retrieval was performed by incubating tissues in either the softening buffer (20% (weight/volume) sodium dodecyl sulfate (SDS), 100 mM β mercaptoethanol, 25 mM ethylenediaminetetraacetic acid (EDTA) and 0.5% Triton-X in 50 mM Tris at pH 8 at RT for 1 hr (**Fig. 1A**) (**Fig. 2A,B; 3L; 4A,G,M,S; 5A,B; 7A**) or by microwave heating for 1 min in 5mM citric acid buffer, 0.5% Triton-X, pH 6, because it provided improved collagen staining (**Supp. Fig. 4A**) (**Supp. Fig. 5A,B**). Antigen retrieval was then followed by 3 washes in 1x PBS for 5 min each and blocking at 37°C for 30 min with MAXblock blocking buffer (Active Motif, #15252)^6^. Immunostaining was performed by diluting primary antibody in MAXbind Staining buffer (Active Motif, #15253), and incubating tissue samples in the antibody solution at 37°C for 1 hr, at RT for 2.5 hr or at 4°C overnight. The same procedure conditions were applied for secondary antibodies. Primary and secondary antibodies used in this work are listed in **Supp. Table 2**. All pre-expansion stained tissues were immersed in VectaShield mounting media (Vector Laboratories, #H-1000-10) and covered with a No. 1 coverslip prior to imaging (**Fig. 2A,B; 3A-C,L; 4A,G,M,S; 5A,B; 7A; Supp. 5A,B; 7A**)

#### Anchoring and gelation

Anchoring and gelation were performed according to previously published protocols^6,19^, and briefly summarized below. Acryloyl-X (a.k.a. 6-((acryloyl)amino)hexanoic acid, succinimidyl ester, here abbreviated AcX, Thermo Fisher Scientific, #A20770) powder was dissolved in anhydrous dimethyl sulfoxide at a concentration of 10 mg/ml and stored in aliquots in a desiccated environment at −20°C. Tissues underwent anchoring by incubation with AcX at a concentration of 0.1 mg/ml in 1x PBS with 0.5% Triton-X, at 4°C for 30 min, followed by 1.5 hrs at 37°C, and then x3 washes with 1x PBS at RT for 5 min each. Next, a monomer solution composed of 1× PBS, 2 M sodium chloride (NaCl), 8.625% (w/v) sodium acrylate, 2.5% (w/v) acrylamide and 0.10% (w/v) N,N′ methylenebisacrylamide (Sigma-Aldrich) was prepared, aliquoted and stored at −20°C. Gelling solution was prepared by mixing the monomer solution with the following chemicals, in the order shown: (1) 4-hydroxy-2,2,6,6-tetramethylpiperidin-1-oxyl (abbreviated as 4-HT; final concentration, 0.01% (w/v)) as an inhibitor of gelation, (2) tetramethylethylenediamine (abbreviated as TEMED; final concentration, 0.2% (w/v)) as an accelerator of gelation, and (3) ammonium persulfate (abbreviated as APS; final concentration, 0.2% (w/v)) as an initiator of gelation. Tissue sections on glass slides were covered with gelling solution, and then a gel chamber was constructed by first placing two No. 1.5 square coverslips (22 mm x 22 mm) as spacers, one at each end of the glass slide and flanking the tissue section in the middle; then, a rectangular coverslip is placed on top of spacers, to enclose the gel chamber, in which the tissue sample is fully immersed in the gelling solution and sandwiched by the glass slide and the top coverslip. Samples were first incubated in a humidified atmosphere at 4°C for 30 min, which slows down gelation rate and enables diffusion of solution into tissues and subsequently incubated in a humidified atmosphere at 37°C for 2.5 hrs to complete gelation (**Fig. 1B; Supp. Fig. 1B, 2B, 3B, 4B**).

#### Softening

After gelation, all coverslips were gently removed from the glass slide that carries the gelled tissue. Excessive gel around the tissue sample was trimmed away using a razor blade. Then, tissues were incubated in the softening buffer, which consists of 20% (w/v) SDS, 100 mM β mercaptoethanol, 25 mM EDTA, 0.5% Triton-X, and Tris 50 mM at pH 8, at 37°C for 30 min followed by 1 hr in an autoclave at 121°C, followed by cooling to RT for 30 min (**Fig. 1C; Supp. Fig. 1C; 2D; 3C; 4D**). Tissues were observed to detach from the glass slides during softening, or during subsequent washes with gentle shaking.

#### Decrowding and expansion without post-expansion immunostaining

After softening, tissue underwent either decrowding (**Fig. 1D; Supp. Fig. 1D**) or expansion without post-expansion immunostaining (**Supp. Fig. 3D; 4E**). For decrowding, tissues were washed 5 times with 1x PBS at RT for 3 min each. At this stage, tissues were at a partially expanded state, with ∼2.3x linear expansion factor. For expansion without post-expansion immunostaining, tissues were then additionally washed in deionized water for 3-5 times at RT for 3 min each, to expand the hydrogel-embedded tissue to an expansion factor of ∼4x ^6,15,19^ (**Supp. Fig. 3D; 4D**). A subset of tissue samples was imaged by confocal microscopy at this state, with methods described in the section *Image Acquisition*, to obtain the post-expansion, pre-decrowding staining images **(Fig. 2C,D; 3E-G; 4B,H,N,T; 5C,D; 7C; Supp. Fig. 5A,B; 6E-F; 7A**).

#### Immunostaining post-decrowding

Then tissues underwent decrowding by washing 5 times with 1x PBS at RT for 3 min which results in an expansion factor of ∼2.3x (**Fig. 1D; Supp. Fig. 1D; 2E; 3E**). Then, we prepared antibody solutions by diluting primary antibody in MAXbind Staining buffer (Active Motif, #15253) and performed post-decrowding immunostaining (**Fig. 1E; Supp. Fig. 1E; 2F; 3F**) by incubating tissue samples in the antibody solution at 37°C for 1 hr, at RT for 2.5 hr or at 4°C overnight. The same procedure conditions were applied for secondary antibodies (**Fig. 3I-K,M; 4C,I,O,U; 5E,F; 6; 7D,E; Supp. Fig. 7; 8**). For tissues that underwent both pre-expansion and post-decrowding staining, antibody concentrations and incubation conditions were identical to ensure quantitative comparisons pre- and post-expansion (**Fig. 4; 5; 7**). Immunostained tissues were expanded by washing with deionized water at RT for 3-5 times for 3 min each to ∼4x linear expansion (**Fig. 1E; Supp. Fig. 3F**). We then performed confocal microscopy at this state, with methods described in the following section *Image Acquisition*, to obtain the post-decrowding-staining images (**Fig. 3I-K,M; 4C,I,O,U; 5E,F; 6; 7D,E; Supp. Fig. 7; 8**).

#### Collagenase treatment

For FFPE 5-µm thick samples of human high-grade glioma brain tumor tissues with high degree of extracellular matrix underwent format conversion and pre-expansion immunostaining (**Supp. Fig. 4A**), followed by anchoring and gelation (**Supp. Fig. 4B**). After gelation, all coverslips were gently removed from the glass slide that carried the gelled tissue. Excessive gels around the tissue sample were trimmed away using a razor blade. Tissues that were designated for the collagenase treatment (**Supp. Fig. 4C**) were submerged in collagenase type II (Thermo Fisher Scientific, #17101015) at 1500 U/ml in Hank’s balanced salt solution (HBSS) for 3 hr at 37°C. Next, the collagenases were inactivated by incubating the sample in the softening buffer for 15 min at room temperature (RT) (**Supp. Fig. 4C**). Then, tissues were incubated in fresh softening buffer and underwent subsequent softening steps (**Supp. Fig. 4D**) and expansion without post-expansion immunostaining (**Supp. Fig. 4E**).

#### Antibody stripping and restaining

To enable highly multiplexed imaging, sequential rounds of antibody stripping and post-decrowding staining were performed. For antibody stripping on tissues stained at the post-decrowding state, we incubated tissues in the softening buffer for 2 hr at 70°C. Afterwards, we washed the tissues 5 times with 1x PBS at RT for 3 min each (**Supp. Fig. 1F**). At this stage, tissues had an expansion factor of ∼2.3x. Samples were then expanded and imaged (**Supp. Fig. 7C**) or subsequently immunostained with only secondary antibodies (**Supp. Fig. 7D**) or both primary and secondary antibodies (**Supp. Fig. 7H,J; 8**).

#### Tissue shrinking

We shrunk tissues to a ∼1.3x linear expansion factor by treating with in a high-ionic-strength buffer (1M NaCl + 60 mM MgCl2)^19^ following the softening (**Supp Fig. 3C**) and washing with PBS (which results in an expansion factor of ∼2.3x) (**Fig. 7B**, pre-decrowding) or following post-decrowding staining (**Fig. 3F**) and washing with PBS (which results in an expansion factor of ∼2.3x) (**Fig. 7D**, post-decrowding). Specifically, we washed the tissues 3-5 times with this buffer at RT for 3 min each, until no more tissue shrinkage was observed. We then performed confocal microscopy at this stage, with methods described in the following section *Image Acquisition*, to obtain pre-decrowding (**Fig. 7B**) or post-decrowding staining at shrunken state images (**Fig. 7D**).

### Imaging processing methods

#### Image acquisition

For confocal imaging, we used a spinning disk confocal system (CSU-W1, Yokogawa) on a Nikon Ti-E microscope. The objective lenses that we used include a 40x 1.15 NA water immersion objective (**Fig. 2C,D; 3A-C,E-G,I-K,L-M; 4A-C,G-I,M-O,S-U; 5C-F; 6; 7; Supp. Fig. 6A-C,E-G; 7A-D,H,J; 8**), or 10x 0.20 NA air objective (**Supp. Fig. 5A-D**). The excitation lasers and emission filters that we used to image each fluorescent dye are the following: 405 nm excitation, 450/50 nm emission filter; 488 nm excitation, 525/40 nm emission filter; 561 nm excitation, 607/36 emission filter; 640 nm excitation, 685/40 emission filter. The following acquisition and display settings apply to all images shown in this study, unless otherwise specified: (1) within the same experiment (as grouped by figures and described in the Results and Figure Legends), all images were obtained with the same laser power, camera settings, and objective lens. (2) For all image display in all figures except **Fig. 4**, brightness and contrast settings were first individually set by the automated adjustment function in ImageJ, and then manually adjusted (raising the minimum-intensity threshold and lowering the maximum-intensity threshold) to improve contrast for features of interest. For image display of **Fig. 4** and **Supp. Fig. 7H**, brightness and contrast settings of images were adjusted so that 1% of the pixels were saturated. None of these changes in the brightness and contrast settings, throughout the entire study, affects the downstream quantitative analysis of fluorescent intensities, which were always applied on raw images, as specified in Results and captions.

For super-resolution structured illumination microscopy (SR-SIM) of samples in the pre-expansion state, for isotropy analyses (**Fig. 2A,B**) and comparative analyses (**Fig. 5A, B**), we used a Deltavision OMX Blaze (GE Healthcare) SR-SIM microscope with a 100x 1.40 NA (Olympus) oil objective to acquire the images.

#### Distortion quantification

We quantified sample distortion based on the same analysis employed in other ExM publications^6,15,19,33,34^, using the custom algorithm developed in MATLAB. In this analysis, we performed B-spline-based non-rigid registration between a pair of images to derive a distortion vector field. We then computed the root-mean-square (RMS) errors on feature measurements in the vector field. This analysis was performed on registrations from pre-expansion and post-expansion images (**Fig. 2E, F, Supp. Fig. 5E; 7K**).

#### Image registration between pre-expansion and post-expansion images

To register a post-expansion image to a given pre-expansion image of the same sample, we first took the entire post-expansion image stack (which thus contained the axial plane corresponding to the entirety of the pre-expansion image), and computed a stack of sum-intensity z-projection images, each of which was a sum of 4 consecutive images in the z-stack (and this moving window of 4 images ran through the entire raw stack, by increments of 1 image in the raw stack; for example, z-projection image #1 was made from raw images #1 - #4, z-projection image #2 was from raw images #2 - #5, and so forth). This procedure ensured that each post-expansion z-projection image covered a similar optical section in biological units given the 4-fold linear expansion of the post-expansion sample. Then, we searched within the stack of post-expansion images (acquired through the entire depth of the tissue sample), for the post-expansion z-projection image that corresponded best to the axial plane of the selected pre-expansion image; by applying a scale-invariant feature transform (SIFT) algorithm followed by random-sample-consensus (RANSAC) with a published, open-source MATLAB package^102^, which registered the pre-expansion image to every post-expansion z-projection image based on program-generated features on these images, and identified the post-expansion z-projection image with the greatest number of matching features. Afterwards, we performed a rigid transformation to rigidly register the pre-expansion image and the post-expansion z-projection image that had the greatest number of matching features (**Fig. 2A,C and B,D; 3A-C and E-G; 3L,M; 4A,B and G,H and M, N and S,T; 5A,C and B,D; Supp. Fig. 5 A,C and B,D; Supp Fig. 6A-C and E-G**).

#### Image registration between post-expansion images and post-decrowding ∼4x expanded Images

To register a post-decrowding image to a given post-expansion image of the same sample, we first took the entire post-decrowding image stack which thus contained the axial plane corresponding to the entirety of the pre-expansion image), and computed a stack of sum-intensity z-projection images, each of which was a sum of 4 consecutive images in the z-stack (and this moving window of 4 images rans through the entire raw stack, by increments of 1 image in the raw stack; for example, z-projection image #1 was made from raw images #1 - #4, z-projection image #2 was from raw images #2 - #5, and so forth). This procedure ensured that each post-decrowding z-projection image covered a similar optical section compared to a given post-expansion z-projection image. Then, we searched within the stack of post-decrowding images (acquired through the entire depth of the tissue sample), for the post-decrowding z-projection image that corresponded best to the axial plane of the matched post-expansion z-projection image; by applying a SIFT-RANSAC algorithm, which identified the post-decrowding z-projection image with the greatest number of matching features to the matched post-expansion z-projection image above. Afterwards, we performed a rigid transformation to rigidly register the matched post-expansion z-projection image and the matched post-decrowding z-projection image (**Fig. 3E-G and I-K; 4B,C and H,I and N,O and T,U; 5C,E and D,F**).

To register other images which were only in ∼4x-expanded states, (**Supp. Fig. 7A-D, H, J; Supp. Fig. 8**) we first selected a sum-intensity z-projection image centered at approximately the mid-z-axial plane image of an image stack, which we assigned as the initial state z-projection image (see **Supp. Fig. 7A**, in which the post-expansion (not stained) z-projection image was the initial state image). Then, each of the subsequent state image stacks (the stacks following after the initial state) was individually registered to the image at the initial state using the workflow from above to find the z-projection image with the greatest number of matching features to the initial z-projection image using DAPI (DAPI not shown) (see **Supp. Fig. 7B-D,** in which post-expansion (stained, stripped x2 hrs, or 2ry antibody only stained) z-projection images were the subsequent state images after the initial state). All registrations were performed as described above: first by applying the SIFT-RANSAC algorithm to identify the corresponding z-projection image from the subsequent stack with the highest number of matching features, and then by performing a rigid transformation to rigidly register the subsequent state z-projection images to the initial state z-projection image.

#### Image Registration between Pre-expansion, Pre-decrowding, Post-decrowding 1x state and ∼4x Expanded Images

To register the five images in **Fig. 7A-E**,, which were at either the pre-expansion, shrunken or ∼4x-expanded states, we performed rigid registrations in the following order using the SIFT-RANSAC algorithm described above: the ∼4x-expanded pre-decrowding-staining z-projection image (**Fig. 7C**) was registered to the pre-expansion single z-slice image (**Fig. 7A**); the pre-decrowding-staining shrunken state single slice image (**Fig. 7B**) was registered to the registered ∼4x-expanded pre-decrowding-staining z-projection image (**Fig. 7C**); the ∼4x-expanded post-decrowding-staining image (**Fig. 7E**) was registered to the ∼4x-expanded pre-decrowding-staining z-projection image (**Fig. 7C**); and then the post-decrowding-staining shrunken state image (**Fig. 7D**) was registered to the ∼4x-expanded post-decrowding-staining z-projection image (**Fig. 7E**).

#### Quantification of Lipofuscin Autofluorescence Removal

To quantify the autofluorescence intensity of lipofuscin between the pre-expansion (not stained) state (**Fig. 3A-C; Supp. Fig. 6A-C**) and the post-expansion (not stained) state (**Fig. 3E-G; Supp. Fig. 6E-G**), we generated masks, and quantified fluorescence after applying them to selected ROIs.

Generation of masks: Autofluorescence from lipofuscin was regarded as the signal in this analysis. Since the autofluorescence was observed in all 3 fluorescent channels imaged (**Fig. 3A-C; Supp. Fig. 6A-C**), we used the 488 nm excitation (abbreviated as “ex”)/525 nm emission (abbreviated as “em”) channel (**Fig. 3A; Supp. Fig. 6A**) as a representative channel and performed segmentation of the lipofuscin aggregates in images from that channel. Pre-expansion images of the 488ex/525em channel were segmented into signal-positive regions (whose pixels were assigned to the signal mask) and signal-negative regions (i.e. all other pixels, which were assigned to the background mask), by manually setting a threshold intensity value, such that the regions whose intensity values were greater than the threshold (thus with sufficiently bright autofluorescence) completely covered the lipofuscin aggregates, by manual inspection. All pixels whose values were greater than the threshold were assigned to the signal mask, and all others were assigned to the background mask.

Selection of regions of interest (ROIs) and fluorescence quantification: For each of the reported mean fluorescence intensities (**Fig. 3D, H; Supp. Fig. 6D, H**), we selected 5 signal and 3 background ROIs per field of view. We imaged 3 fields of view for each sample. We evaluated 3 tissue samples each from a different patient. For each signal ROI, the reported fluorescence intensity was computed from the mean fluorescence intensity value across the entire signal ROI, in either the pre-expansion images (**Fig. 3D; Supp. Fig. 6D**) or the post-expansion images (**Fig. 3H; Supp. Fig. 6H**). ROIs have a dimension of 5×5 pixels (corresponds to 0.2 microns in biological units). The signal and background ROIs were selected based on the following criteria:

Lipofuscin and Background (**Fig. 3D-F**)

*Signal ROIs*

- Lipofuscin-positive ROI, in normal cortex tissue: ROIs that fit entirely within the lipofuscin-signal mask
*Background ROIs*

- Background ROI, in normal cortex tissue: ROIs that fit entirely within the background mask that were at least one ROI width away from the pixels that were positive for the lipofuscin-signal mask.
*Mean Fluorescence Intensity Calculations*

- We calculated the mean fluorescence intensities in the 488ex/525em channel (**Fig. 3D, H**; **Supp. Fig. 6D, H**; cyan), 561ex/607em channel (**Fig. 3D, H**; **Supp. Fig. 6D, H**; yellow), and 640ex/685em channel (**Fig. 3D, H**; **Supp. Fig. 6D, H**; magenta) for the lipofuscin-positive ROIs (**Fig. 3D, H**; **Supp. Fig. 6D, H**; left) and background ROIs (**Fig. 3D, H**; **Supp. Fig. 6D, H**; right), respectively, in both the pre-expansion (**Fig. 3D; Supp. Fig. 6D**) and post-expansion images (**Fig. 3H; Supp. Fig. 6H**).
- Statistical analysis: We averaged the mean fluorescence value of all 5 signal (and of all 3 background) ROIs in each field of view and given 4 samples and 3 field of view per sample, a total of 12 mean signal and 12 mean background fluorescence intensity values for the lipofuscin-positive signal and background ROIs for each channel in the pre-expansion (**Fig. 3D; Supp. Fig. 6D**) and post-expansion (**Fig. 3H; Supp. Fig. 6H**) images. Box plot: individual values (open circles), median (middle line), mean (dotted line), first and third quartiles (lower and upper box boundaries), lower and upper raw values (whiskers). We then applied a 2-tailed paired t-test (non-Bonferroni corrected) to lipofuscin vs. background, for pre-expansion mean fluorescence intensities for each spectral channel and also separately for post-expansion mean fluorescence intensities for each spectral channel, with p < 0.05 considered statistically significant.

#### Fluorescence quantification for protein decrowding

To quantify post-decrowding staining, pixel intensity values were compared between post-expansion (not restained) (**Fig. 4B, H, N, T)** and post-expansion (restained) images (**Fig. 4C, I, O, U**), as expansion factor strongly affects pixel intensity values. Quantitative analysis was conducted as follows.

Generation of the signal mask: We constructed a binary image “signal” mask, for each stain, that corresponded to positive pixels (i.e., above a manually selected threshold; we were not blinded to condition) for a given stain in both pre-expansion and post-expansion staining images. All images were segmented into signal-positive pixels according to the threshold intensity value for each image. All pixels whose values were greater than the thresholds in both post-expansion (no restained) and post-expansion (restained) images were assigned to the signal mask.

Generation of background mask: Because post-expansion (restained) images revealed additional structures not visible in pre-expansion or post-expansion (not restained) images, we created a second “background” mask for each stain in each of the pre-expansion and post-expansion (restained) images, corresponding to negative pixels that were below the threshold used for the signal mask. Then, a “doubly negative” background mask was constructed for each stain, corresponding to the pixels that were negative in both the pre-expansion and post expansion (restained) image background masks for that stain.

Selection of regions of interest (ROIs) and fluorescence quantification: For each of the reported mean fluorescence intensities (**Fig. 4D-F; 4J-L; 4P-R; 4V-X**), we evaluated 3 tissue samples, each from a different patient, with 3 fields of view for each sample, and selected 5 signal and 3 background ROIs per field of view. For each signal ROI, the reported fluorescence intensity was computed from the mean fluorescence intensity value across the entire signal ROI, in either the post-expansion (not restained) staining images (**Fig. 4B, H, N, T**) or the post-expansion (restained) images (**Fig. 4C, I, O, U**), which were both ∼4x-expanded sample states. ROIs corresponded to 0.2 microns in biological units (or 5×5 pixels). The signal and background ROIs were selected based on the following criteria, manually selected, without blinding to condition:

MAP2 and GFAP (**Fig. 4D-F**)

*Signal ROIs*

- MAP2-positive ROIs, in normal hippocampus tissue: ROIs that fit entirely within the MAP2-signal mask (i.e., all 25 pixels were signal positive) that were at least one ROI width away from the GFAP-signal mask
- GFAP-positive ROIs, in normal hippocampus tissue: ROIs that fit entirely within the GFAP-signal mask that were at least one ROI width away from the MAP2-signal mask
*Background, Doubly Negative ROIs*

- MAP2 and GFAP doubly negative ROIs, in normal hippocampus tissue: ROIs that fit entirely within the doubly negative mask that were at least one ROI width away from the pixels that were positive for either the MAP2- or GFAP-signal masks.
*Mean Fluorescence Intensity Calculations*

- We calculated the mean fluorescence intensities of MAP2-positive ROIs, in the MAP2 channel (**Fig. 4D**, left, cyan) and in the GFAP channel (**Fig. 4D**, right, magenta) in the post-expansion (not restained, NR) and post-expansion (restained, R) images. Box plot: individual values (open circles), median (middle line), mean (dotted line), first and third quartiles (lower and upper box boundaries), lower and upper raw values (whiskers). We did the same procedure for the GFAP-positive ROIs (**Fig. 4E**) and the doubly negative ROIs (**Fig. 4F**).
GFAP and α-SMA (**Fig. 4J-L**)

*Signal ROIs*

- GFAP-positive ROIs, in high-grade glioma tissue: ROIs that fit entirely within the GFAP-signal mask that were at least one ROI width away from the α SMA-signal mask
- α--SMA-positive ROIs, in high-grade glioma tissue: ROIs that fit entirely within the α-SMA-signal mask that were at least one ROI width away from the GFAP-signal mask
*Background, Doubly Negative ROIs*

- GFAP and α-SMA doubly negative ROIs, in high-grade glioma tissue: ROIs that fit entirely within the doubly negative mask that were at least one ROI width away from the pixels that were positive for either the GFAP- or α-SMA-signal masks.
*Mean Fluorescence Intensity Calculations*

- We calculated the mean fluorescence intensities of GFAP-positive ROIs, in the GFAP channel (**Fig. 4J**, left, cyan) and in the α SMA channel (**Fig. 4J**, right, magenta) in the post-expansion (not restained, NR) and post-expansion (restained, R) images. Box plot: individual values (open circles), median (middle line), mean (dotted line), first and third quartiles (lower and upper box boundaries), lower and upper raw values (whiskers). We did the same procedure for the α SMA-positive ROIs (**Fig. 4K**) and the doubly negative ROIs (**Fig. 4L**)
Vimentin and α-SMA (**Fig. 4P-R)**

*Signal ROIs*

- Vimentin-positive ROIs, in high-grade glioma tissue: ROIs that fit entirely within the vimentin-signal mask that were at least one ROI width away from the α SMA-signal mask
- α-SMA-positive ROIs, in high-grade glioma tissue: ROIs that fit entirely within the α-SMA-signal mask that appeared to have as low as possible an amount of vimentin, yet as noted in the results, there was a high likelihood, that some vimentin-positive signal pixels were found in α-SMA ROIs, unlike the methods noted above in which the α-SMA-positive ROIs were at least one ROI width away from the GFAP-signal mask.
*Background, Doubly Negative ROIs*

- Vimentin and α SMA doubly negative ROIs, in high-grade glioma tissue: ROIs that fit entirely within the doubly negative mask that were at least one ROI width away from the pixels that were positive for either the vimentin- or α SMA-signal masks.
*Mean Fluorescence Intensity Calculations*

- We calculated the mean fluorescence intensities of vimentin-positive ROIs, in the vimentin channel (**Fig. 4P**, left, cyan) and in the α-SMA channel (**Fig. 4P**, right, magenta) in the post-expansion (not restained, NR) and post-expansion (restained, R) images. Box plot: individual values (open circles), median (middle line), mean (dotted line), first and third quartiles (lower and upper box boundaries), lower and upper raw values (whiskers). We did the same procedure for the α SMA-positive ROIs (**Fig. 4Q**) and the doubly negative ROIs (**Fig. 4R**)
Iba1 and GFAP (**Fig. 4V-X**)

*Signal ROIs*

- Iba1-positive ROIs, in low-grade glioma tissue: ROIs that fit entirely within the Iba1-signal mask that were at least one ROI width away from the GFAP-signal mask
- GFAP-positive ROIs, in low-grade glioma tissue: ROIs that fit entirely within the GFAP-signal mask that were at least one ROI width away from the Iba1-signal mask
*Background ROIs*

- Iba1 and GFAP doubly negative ROIs, in low-grade glioma tissue: ROIs that fit entirely within the doubly negative mask that were at least one ROI width away from the pixels that were positive for either the Iba1- or GFAP-signal masks.
*Mean Fluorescence Intensity Calculations*

- We calculated the mean fluorescence intensities of Iba1-positive ROIs, in the Iba1 channel (**Fig. 4V**, left, cyan) and in the GFAP channel (**Fig. 4V**, right, magenta) in the post-expansion (not restained, NR) and post-expansion (restained, R) images. Box plot: individual values (open circles), median (middle line), mean (dotted line), first and third quartiles (lower and upper box boundaries), lower and upper raw values (whiskers). We did the same procedure for the GFAP-positive ROIs (**Fig. 4W**) and the doubly negative ROIs (**Fig. 4X**)

Statistical analysis: We averaged the mean fluorescence value of all 5 signal ROIs and all 3 doubly negative ROIs in each field of view. With 3 samples and 3 fields of view per sample, a total of 9 mean signal fluorescence intensity values for the MAP2-positive signal mask ROIs for pre- and post-decrowding images in the MAP2-channel (**Fig. 4D**, left, cyan) and GFAP-channel (**Fig. 4D**, right, magenta), and 9 mean doubly negative fluorescence intensity values for the MAP2/GFAP images were calculated. We performed the same calculations for GFAP/α MA (**Fig. 4J**), vimentin/α MA (**Fig. 4P**), and Iba1/GFAP (**Fig. 4V**). We then applied a 2-tailed paired t-test, (non-Bonferroni corrected) to each post-expansion (not restained) and post-expansion (restained) set of 9 averaged values (**Fig. 4D-F, J-L, P-R, and W-X**), with p < 0.05 considered statistically significant.

#### Quantification of fluorescence co-localization of vimentin, Iba1, and GFAP in in low-grade gliomas

To quantify the co-localization of the markers vimentin, GFAP and Iba1, in the pre-expansion-stained state (**Fig. 7F-H**, gray boxplots) and the post-decrowding staining at shrunken state (**Fig. 7F-H**, white boxplots) images, we performed the following analysis.

Generation of nuclei masks: We constructed a binary image “nuclei” mask, corresponding to pixels positive for nuclei (DAPI) in pre-expansion images. Pre-expansion images were each segmented into nuclei-positive pixels using a publicly-available automated, deep learning based segmentation method called Cellpose (www.cellpose.org)^103^. Gray scale (DAPI channel) images of DAPI-stained pre-expansion images were uploaded to the Cellpose algorithm, which provided an output of segmented nuclei. From the segmented nuclei, we extracted the edges of each nucleus, as well as the centroid, and calculated the total number of nuclei per pre-expansion image.

Generation of signal mask: We constructed binary image “signal” masks for each stain, corresponding to pixels that were positive for that stain in pre-expansion or post-decrowding stained at shrunken state images. Pre-expansion and post-decrowding staining images for each stain were each segmented into signal-positive pixels using an automated Otsu’s segmentation algorithm^104^ in Matlab to calculate a threshold intensity value for each image. All pixels whose values were greater than the automatically determined threshold in the pre-expansion image were assigned to the pre-expansion signal mask, and all pixels whose values were greater than the automatically determined threshold in the post-decrowding staining image were assigned to the post-decrowding signal mask. We calculated this for each individual stain.

#### Fluorescence quantification

##### Percent of positive pixels

Next, we quantified the percent positive pixels among all pixels in each field of view within each stain (vimentin, V; Iba1, I; GFAP, G), or combination of stains (Iba1 and vimentin, I&V; vimentin and GFAP, V&G; Iba1 and GFAP, I&G) (**Fig. 7F**). First, for each individual stain we counted the number of pixels in the signal mask that were positive for that stain. Then we counted the total number of pixels (positive or not) in the field of view. We then calculated the percent positive pixels for that stain by dividing the number of positive signal pixels by the total number of pixels in the field of view and multiplying by 100. Next, for each combination of stains, we counted the number of pixels that were “doubly positive” for both stains using the signal masks for each individual stain. We then calculated the percent of “doubly positive” pixels for that combination of stains by dividing the number of “doubly positive” signal pixels over the total number of pixels in the field of view and multiplying by 100. We performed this pixel quantification method for the pre-expansion images (**Fig. 7F,** gray colored bars) and then also for the post-decrowding images (**Fig. 7F,** black colored bars).

##### Number of positive cells

Next, we quantified the total number of positive cells in each field of view for each stain (vimentin, V; Iba1, I; GFAP, G), or combination of stains (Iba1 and vimentin, I&V; vimentin and GFAP, V&G; Iba1 and GFAP, I&G) (**Fig. 7G**). First, for each individual stain we created an image overlay which consisted of the signal mask for a single stain (such as a vimentin-positive signal mask displayed in white) and of the nuclei mask (which displayed the centroids in red and the nuclei boundary in green).

We observed cell nuclei and considered a nucleus “positive” for a stain when at least 25% of the linear surface of the nuclear boundary (nuclei boundary displayed in green; nuclei centroid displayed in red) was surrounded by positive signal pixels such that the sum of the pixels surrounded the nuclear boundary was at least >25% (displayed in white).

Using manual selection via a graphical user interface, cells were considered “positive” for a stain if >25% of the cell nuclei boundary (nuclei boundary displayed in green; nuclei centroid displayed in red) was in contact or surrounded by the positive signal pixels (displayed in white). To label a cell as positive for a stain, we visually inspected each image overlay and manually selected the nuclei user a graphical user interface. This manual selection was used to calculate the total number of positive cells for each stain (vimentin, V; Iba1, I; GFAP, G) in pre-expansion and post-decrowding images. From the nuclei selected for each individual stain, we then calculated the cells that were “doubly positive” for each combination of stains (Iba1 and vimentin, I&V; vimentin and GFAP, V&G; Iba1 and GFAP, I&G) (**Fig. 7G**).

Next, with the cells counted above, we could then calculate the percent of positive cells with co-localized staining among all cells that were positive for a single type of stain in the field of view. For example, to calculate the percent of cells that were “doubly positive” (that co-localized) for GFAP and vimentin (**Fig. 7G**, G&V) among all cells that were positive for GFAP (**Fig. 7G**, G), we divided the number of cells calculated above that were “doubly positive” for GFAP and vimentin (G&V) by the number of cells positive for GFAP (G) x 100. We performed the same analysis using the number of cells calculate above for the other combinations of stains in **Fig. 7H** (G&V/V; I&V/I; I&V/V; I&G/I; I&G/G)

Statistical analysis: We calculated the positive pixels, “doubly positive” pixels, or number of positive or “doubly positive” cells in each field of view, and given 3 samples and 2 fields of view per sample, a total of 6 values for the number of positive pixels, “doubly positive” pixels, or number of positive or “doubly positive” cells for each stain (or combination of stains) in the pre- and post-decrowding staining images. Box plot: individual values (open circles), median (middle line), mean (dotted line), first and third quartiles (lower and upper box boundaries), lower and upper raw values (whiskers), used throughout the graphs of this figure. We then applied a 2-tailed paired t-test (non-Bonferroni corrected) for the set of values across all 6 values comparing the pre- and post-decrowding staining images, p < 0.05 considered statistically significant.

#### Quantification of fluorescence signal change with antibody stripping

To quantify the fluorescent intensity of post-decrowding immunostaining at different stages of the antibody stripping protocol (**Supp. Fig. 7A-D**), we performed the following analysis.

Selection of ROIs and fluorescence quantification: For each stain (histone, **Supp. Fig. 7E**; vimentin, **Supp. Fig. 7F**; GFAP, **Supp. Fig. 7G**), we selected 10 signal ROIs on the post-decrowding-staining images (i.e., **Fig. 7B**, because this is the state where we can clearly identify positive regions for each stain) from each of the 4 tissue samples derived from 2 patients (2 samples per patient), for a total of 40 signal ROIs per stain. ROIs have a dimension of 15×15 pixels (corresponds to 0.6 microns in biological units) and were selected on regions with relatively high fluorescence intensities for each stain, to be rigorous about confronting any residual staining, by manual inspection. For each signal ROI, we computed the average intensity value across the entire signal ROI, in the post-expansion state (not stained), post-expansion (stained), post-expansion states (stripped for 1 hr), post-expansion state (stripped for 2 hrs), and post-expansion (2ry antibody only stained). The population statistics of these average intensities were reported in **Supp. Fig. 7E-G** for the three analyzed stains.

Statistical analysis: For the fluorescent intensity measurements, we first group the 40 measurements by their tissue sample (n = 4 samples, with 10 ROIs each), using the average of the 10 measurements in each tissue sample as the representative quantity. We pre-grouped the measurements by tissue sample. We then performed a 2-tailed paired t-test (non-Bonferroni corrected) analysis, between the representative quantities obtained from the post-decrowding-staining image, and images acquired at the other states, p < 0.05 considered statistically significant.

#### Quantification of fluorescence signal change with multiple rounds of immunostaining

To quantify the fluorescent intensity of post-decrowding immunostaining for vimentin, in consecutive rounds of antibody stripping and re-staining (**Supp. Fig. 7I**), we performed the following analysis.

Selection of ROIs and fluorescence quantification: From the image corresponding to the first round of immunostaining (“Round 1” in **Supp. Fig. 7H**), we selected 15 signal ROIs across 2 fields of view from 3 tissue samples. ROIs have a dimension of 0.6 microns, or 15×15 pixels, and were selected as regions with relatively high fluorescence intensities for vimentin, by manual inspection. For each signal ROI, we computed the average intensity value across the entire signal ROI, and we did this for each image acquired at each of the four rounds of immunostaining (Rounds 1 and 4 shown in **Supp. Fig. 7H**). The population statistics of these average intensities were reported in **Supp. Fig. 7I**.

Statistical analysis: For the fluorescent intensity measurements, we calculated the average of the 15 ROI measurements in each field of view for each tissue sample to calculated a representative quantity for each field of view for a total of 6 values for each round. We pre-grouped the measurements by tissue sample. We then performed a 2-tailed paired t-test (non-Bonferroni corrected) analysis between each of the four different rounds of immunostaining, p< 0.05 considered statistically significant.

We then performed a 2-tailed paired t-test (non-Bonferroni corrected) analysis, between the representative quantities obtained for each round of staining, p < 0.05 considered statistically significant.

## Acknowledgments

The study was supported by Lisa Yang, HHMI, John Doerr, Open Philanthropy, the Bill & Melinda Gates Foundation, the Koch Institute Frontier Research Program, NIH 1R01MH123403, NIH R56NS117465, NIH 1R01MH123977, NIH 1R56AG069192, NIH R01MH124606, and NIH 1R01EB024261 (ESB) and the Neurosurgery Research and Education Foundation (PAV).

## Author contributions

PAV, EAC, ESB co-designed the study. PAV performed the human tissue experiments. PAV, BA, DB performed the animal tissue experiments. PAV, CCY performed data analysis. PAV, CCY, JA, YZ, JSB, DB, BA, MSV, KS, EAC, ESB contributed to interpretations of the results. PAV, CCY, JA, JSB, MSV, ESB wrote the manuscript. All authors critically reviewed the manuscript. PAV, EAC, ESB had full access to data and take responsibility for integrity and accuracy.

## Competing interests statement

PAV, YZ, and ESB have filed for patent protection on a subset of the technologies described. CCY is a co-inventor on two different expansion microscopy technologies. JDB has an equity position in Avidea Technologies, Inc., which is commercializing polymer-based drug delivery technologies for immunotherapeutic applications. JDB has an equity position in Treovir LLC, an oHSV clinical stage company and is a member of the POCKiT Diagnostics Board of Scientific Advisors. ESB is cofounder of a company to help with commercial applications of expansion microscopy. JY, JA, DB, BA, JSB, MSV, KS, and EAC declare no competing interests associated with this manuscript.

## Data and materials availability

The dExPath protocol will be posted online at http://expansionmicroscopy.org at time of publication. Data are available upon reasonable request to the corresponding authors of the paper

## Code availability

Code used in this study are available upon reasonable request to the corresponding authors of the paper

## SUPPLEMENTAL MATERIALS

**Supp. Fig. 1.**
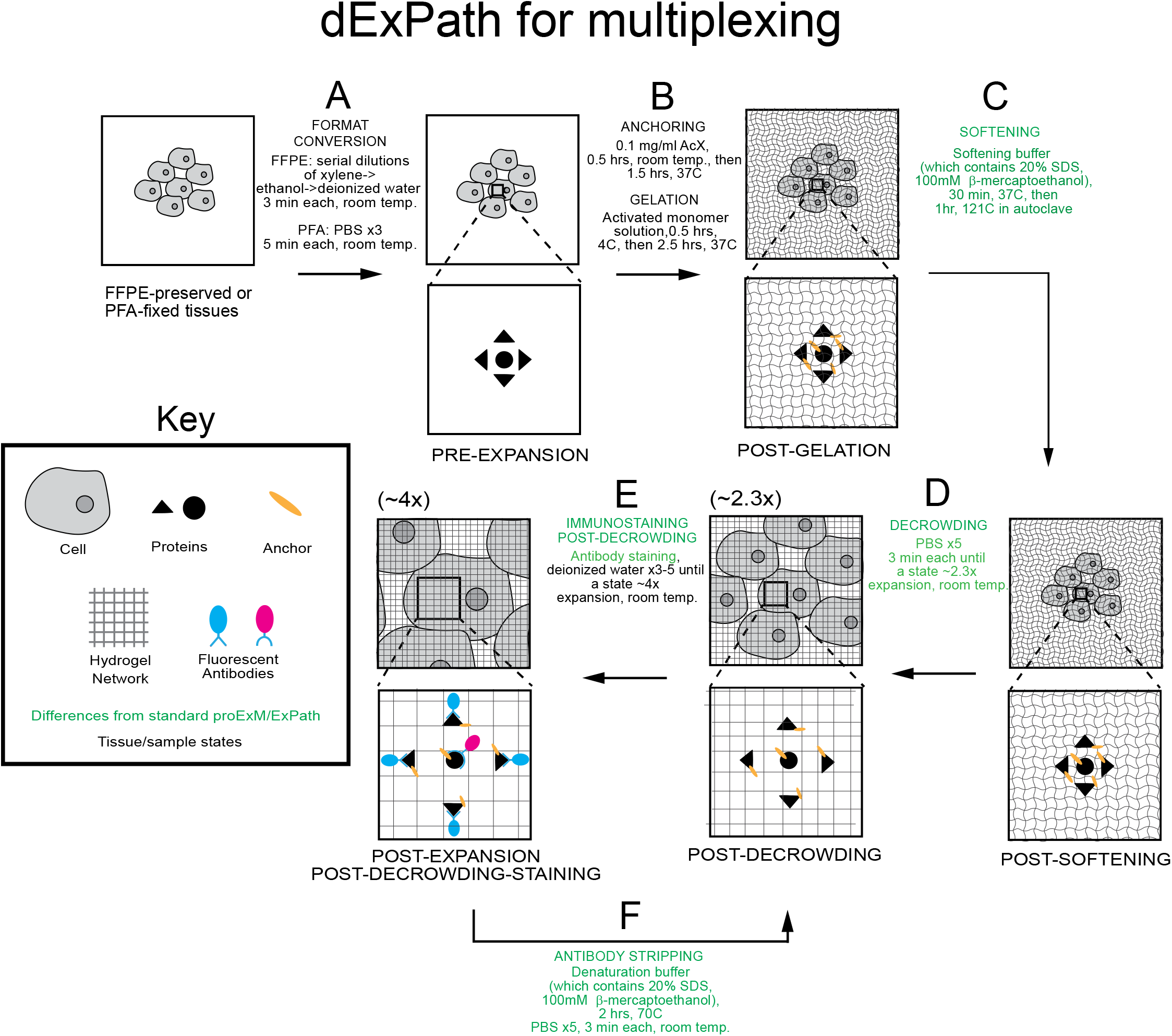
dExPath for highly multiplexed post-expansion immunostaining of formaldehyde-fixed specimens. (A-E) Workflow for expanding FFPE (formalin-fixed paraffin-embedded), or formaldehyde-fixed, human (or mouse) brain specimens, enabling multiple rounds of sequential post-decrowding immunostaining. Key modifications of published proExM and ExPath protocols are shown in green. PFA, paraformaldehyde; PBS, phosphate buffered saline; AcX, Acryloyl-X; SDS, sodium dodecyl sulfate. For steps after decrowding (D), linear expansion factor of the hydrogel-specimen composite is shown in parentheses above the schematic of the step. (A) Tissue samples undergo conversion into a state compatible with expansion. (B) Tissue samples are treated so that gel-anchorable groups are attached to proteins, then the sample is permeated with an expandable polyacrylate hydrogel. (C) Samples are incubated in a softening buffer to denature, and loosen disulfide bonds and fixation crosslinks between proteins in the sample. (D) Softened samples are washed in a buffer to partially expand them. (E) Samples are stained and then expanded fully by immersion in water. (F) Samples undergo repeated rounds of sequential antibody stripping by incubation in softening buffer to remove antibodies, which shrinks the specimen back to 1x, followed by re-expansion to 2.3x, post-decrowding immunostaining and full expansion (E) to enable highly multiplexed imaging.

**Supp. Fig. 2.**
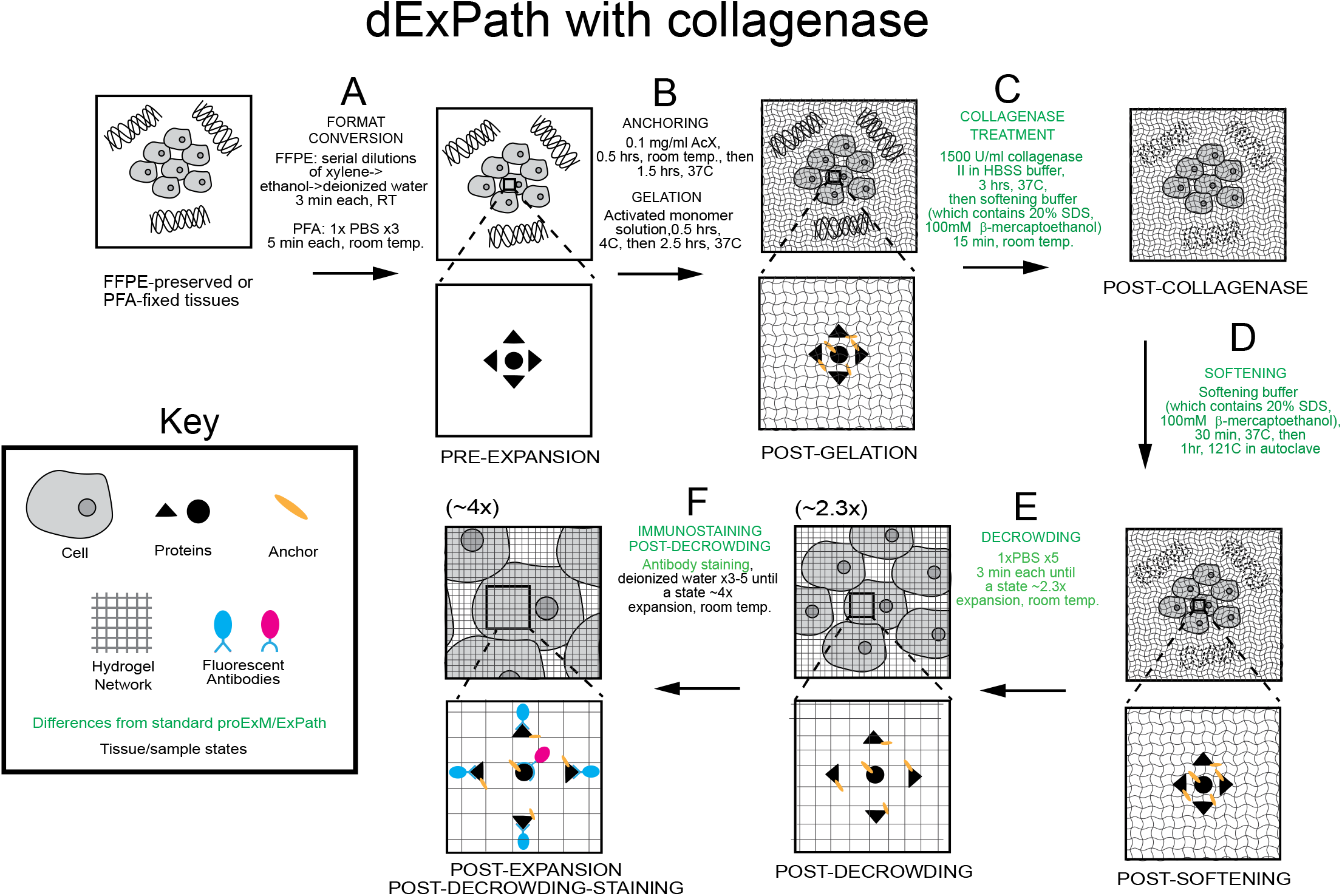
dExPath for post-expansion immunostaining of formaldehyde-fixed specimens with a high degree of extracellular matrix. (A-F) Workflow for expanding FFPE, or formaldehyde-fixed, human (or mouse) brain specimens, enabling post-decrowding immunostaining in human brain tissues with a high degree of extracellular matrix. HBSS, Hank’s balanced salt solution. Key modifications of published proExM and ExPath protocols are shown in green. For steps after decrowding (E), linear expansion factor of the hydrogel-specimen composite is shown in parentheses above the schematic of the step. (A) Tissue samples undergo conversion into a state compatible with expansion. (B) Tissue samples are treated so that gel-anchorable groups are attached to proteins, then the sample is permeated with an expandable polyacrylate hydrogel. (C) Samples are incubated in a buffer containing collagenase. (D) Samples are incubated in a softening buffer to denature, and loosen disulfide bonds and fixation crosslinks between, proteins in the sample. (E) Softened samples are washed in a buffer to partially expand them. (F) Samples are stained and then expanded fully by immersion in water.

**Supp. Fig. 3.**
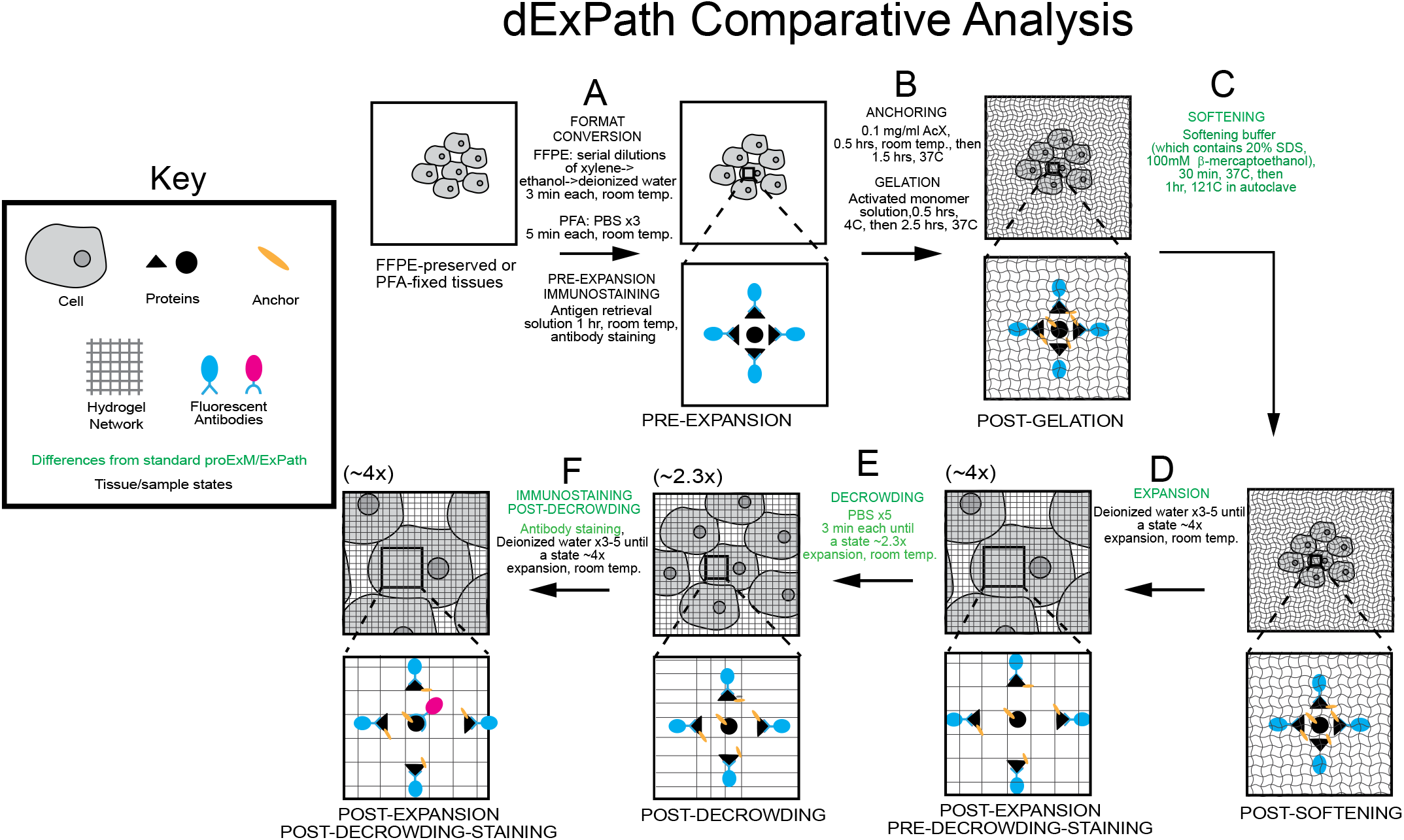
dExPath for post-expansion immunostaining of formaldehyde-fixed specimens that enables comparison of pre- and post-expansion immunostaining. (A-F) Workflow for expanding FFPE, or formaldehyde-fixed, human (or mouse) brain specimens, enabling comparison of pre- and post-decrowding immunostaining. Key modifications of published proExM and ExPath protocols are shown in green. For steps after expansion (D), linear expansion factor of the hydrogel-specimen composite is shown in parentheses above the schematic of the step. (A) Tissue samples undergo conversion into a state compatible with expansion, followed by antigen retrieval and pre-expansion immunostaining. (B) Tissue samples are treated so that gel-anchorable groups are attached to proteins, then the sample is permeated with an expandable polyacrylate hydrogel. (C) Samples are incubated in a softening buffer to denature, and loosen disulfide bonds and fixation crosslinks between, proteins in the sample. (D) Softened samples are then fully expanded for comparative analysis. (E) Expanded samples are converted into a state similar to the decrowded state (∼2.3x) prior to immunostaining. (F) Samples are additionally stained and then expanded fully for comparative analysis.

**Supp Fig 4.**
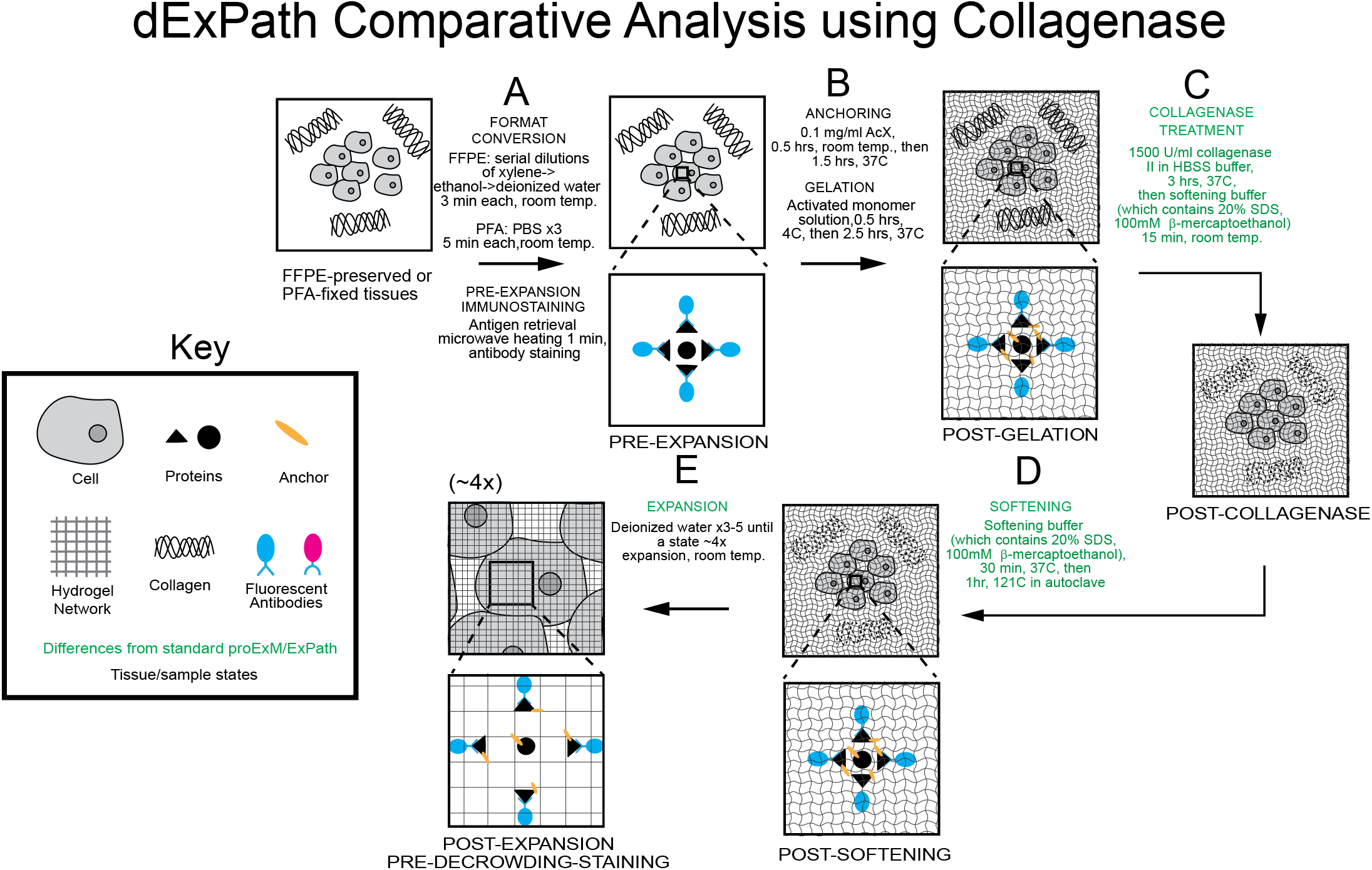
dExPath of formaldehyde-fixed specimens with a high degree of extracellular matrix that enables comparison of pre- and post-expansion tissues. (A-E) Workflow for expanding FFPE, or formaldehyde-fixed, human (or mouse) brain specimens with a high degree of extracellular matrix, enabling comparison of pre- and post-expansion tissues. Key modifications of published proExM and ExPath protocols are shown in green. Following expansion (E), linear expansion factor of the hydrogel-specimen composite is shown in parentheses above the schematic of the step. (A) Tissue samples undergo conversion into a state compatible with expansion, followed by antigen retrieval and pre-expansion immunostaining. (B) Tissue samples are treated so that gel-anchorable groups are attached to proteins, then the sample is permeated with an expandable polyacrylate hydrogel. (C) Samples are incubated in a buffer containing collagenase. (D) Samples are incubated in a softening buffer to denature, and loosen disulfide bonds and fixation crosslinks between proteins in the sample. (E) Softened samples are then fully expanded for comparative analysis.

**Supp. Fig. 5.**
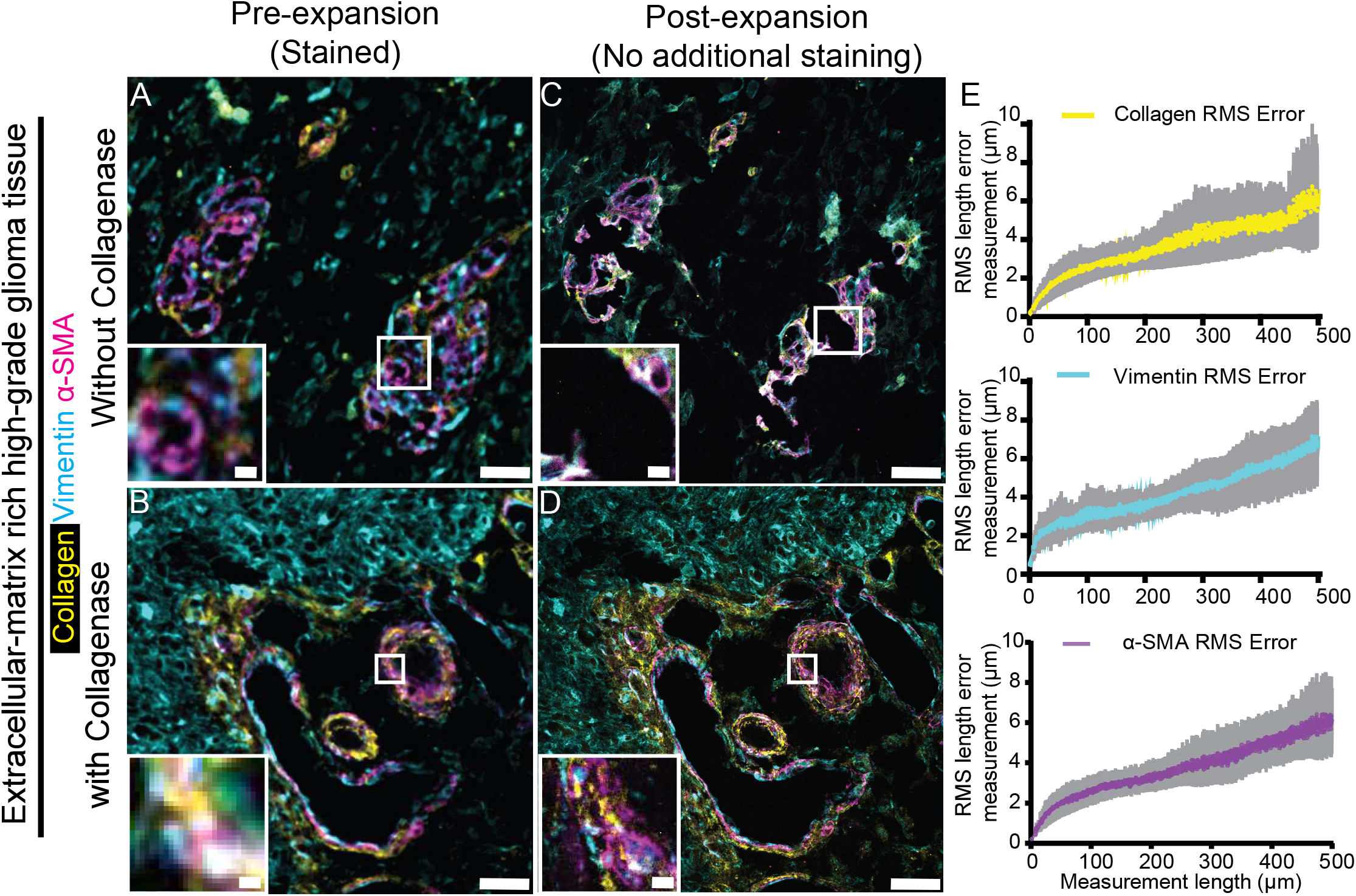
Isotropy of dExPath, with and without collagenase, in human glioma tissues with a high degree of extracellular matrix. (A-B) Representative pre-expansion confocal images of FFPE 5-µm-thick slices of extracellular matrix-rich human high grade-glioma brain tumor tissue (A and B, both n = 3 samples, each from a different patient) which underwent processing as in **Supp. Fig. 4A** (tissue deparaffinization, rehydration, antigen retrieval, and immunostaining)), with immunostaining being for collagen, vimentin, and α SMA, and staining for DAPI (not shown; used for initial rigid alignment). White boxes in (A-B) mark extracellular matrix-rich regions; lower left inset is zoomed-into image of the upper right white box. (C-D) Post-expansion images of the same fields of view as shown in (A-B), respectively. Specifically, samples were treated with anchoring and gelation (as in **Supp. Fig. 4B**), and either no collagenase treatment followed by softening (C), or collagenase treatment followed by softening (D) (as in **Supp. Fig. 4C-D**), and another round of DAPI staining, ∼4x linear expansion (as in **Supp. Fig. 4E**), and imaging with confocal microscopy. White boxes in (C-D), as in (A-B). (E) Root mean square (RMS)-length measurement errors obtained by comparing pre- and post-expansion images for collagenase-treated samples, such as shown in B and D (n = 3 samples, each from a different patient). Line, mean; shaded area, standard deviation. All images are single z-slice confocal images of pre-expansion images (A-B) or sum intensity z-projections of confocal image stacks (C-D), both covering an equivalent tissue depth in biological units. Brightness and contrast settings in images (A-D): first set by the ImageJ auto-scaling function, and then manually adjusted (by raising the minimum-intensity threshold and lowering the maximum-intensity threshold) to improve contrast for stained structures but quantitative analysis in (E) was conducted on raw image data. Scale bars (in biological units): (A-D) outer panel 15 µm; inset, 3 µm. Linear expansion factors: (C-D) 4.0x.

**Supp. Fig. 6.**
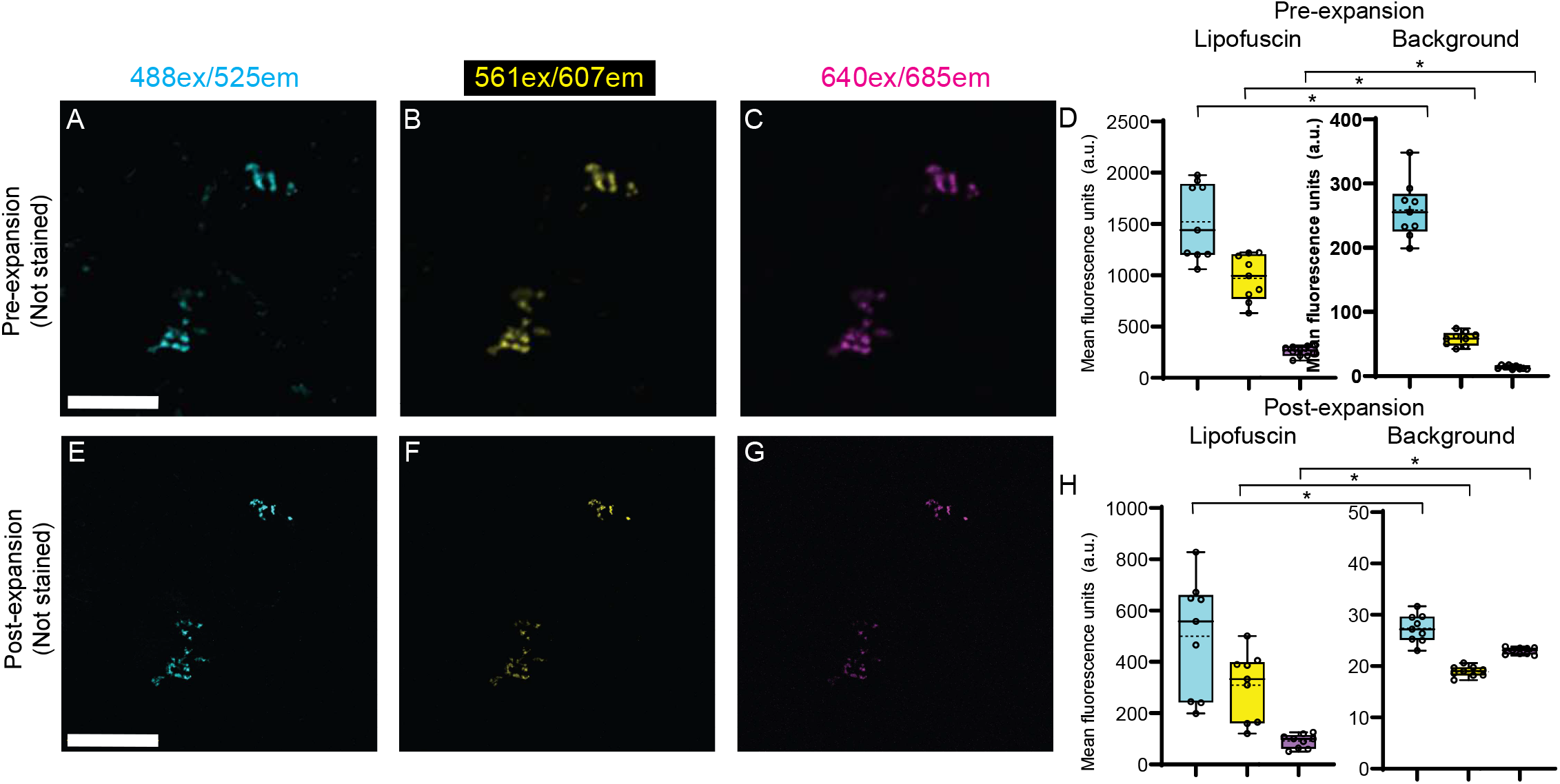
Classical ExPath does not reduce lipofuscin autofluorescence to background levels. (A-C) Representative (n = 3 samples, each from a different patient) pre-expansion confocal images (single z slices) of a neuron of an FFPE 5-µm-thick sample of normal human cortex. The samples underwent format conversion (as in Fig. 1A, tissue deparaffinization and rehydration), and DAPI staining (images not shown in this figure; used for registration across images). Images were acquired for 3 common fluorescent filter settings: (A) 488 nm excitation (abbreviated as “ex”) / 525 nm emission (abbreviated as “em”) channel; (B) 561ex/607em channel; (C) 640ex/685em channel. (D) Mean fluorescence intensities from pre-expansion images, averaged across regions of interest (ROIs) that exhibited prominent lipofuscin (left bar graph), as well as across background ROIs (right bar graph); colors correspond to the colors of A-C (n = 3 tissue samples, each from a different patient). Brightness and contrast settings in images (A-C): first set by the ImageJ auto-scaling function, and then manually adjusted (by raising the minimum-intensity threshold and lowering the maximum-intensity threshold) to improve contrast for lipofuscin but quantitative analysis in (D) was conducted on raw image data. Box plot: individual values (open circles; 3 measurements were acquired from each patient), median (middle line), mean (dotted line), first and third quartiles (lower and upper box boundaries), lower and upper raw values (whiskers). Statistical testing: 2-tailed paired t-test (non-Bonferroni corrected) was applied to lipofuscin vs. background, for pre-expansion mean fluorescence intensities for each spectral channel. *, p < 0.05. (E-G) Post-expansion confocal images after the sample from A-C was additionally treated with anchoring, gelation, digestion with proteinase K, DAPI staining, and ∼4x linear expansion following the proteinase-based ExPath protocol. Sum intensity z-projections of image stacks correspond to the biological thickness of the original slice, taken under identical settings and of the same field of view as A-C and displayed under the same settings. (H) Mean fluorescence intensities, from post-expansion images, averaged across the same lipofuscin (left) and background (right) ROIs used in panel D, for the same samples as panel D. Plots and statistics as in D. Scale bars (in biological units): (A, E) 7 µm; linear expansion factor: (E-G) 4.4x.

**Supp Fig. 7.**
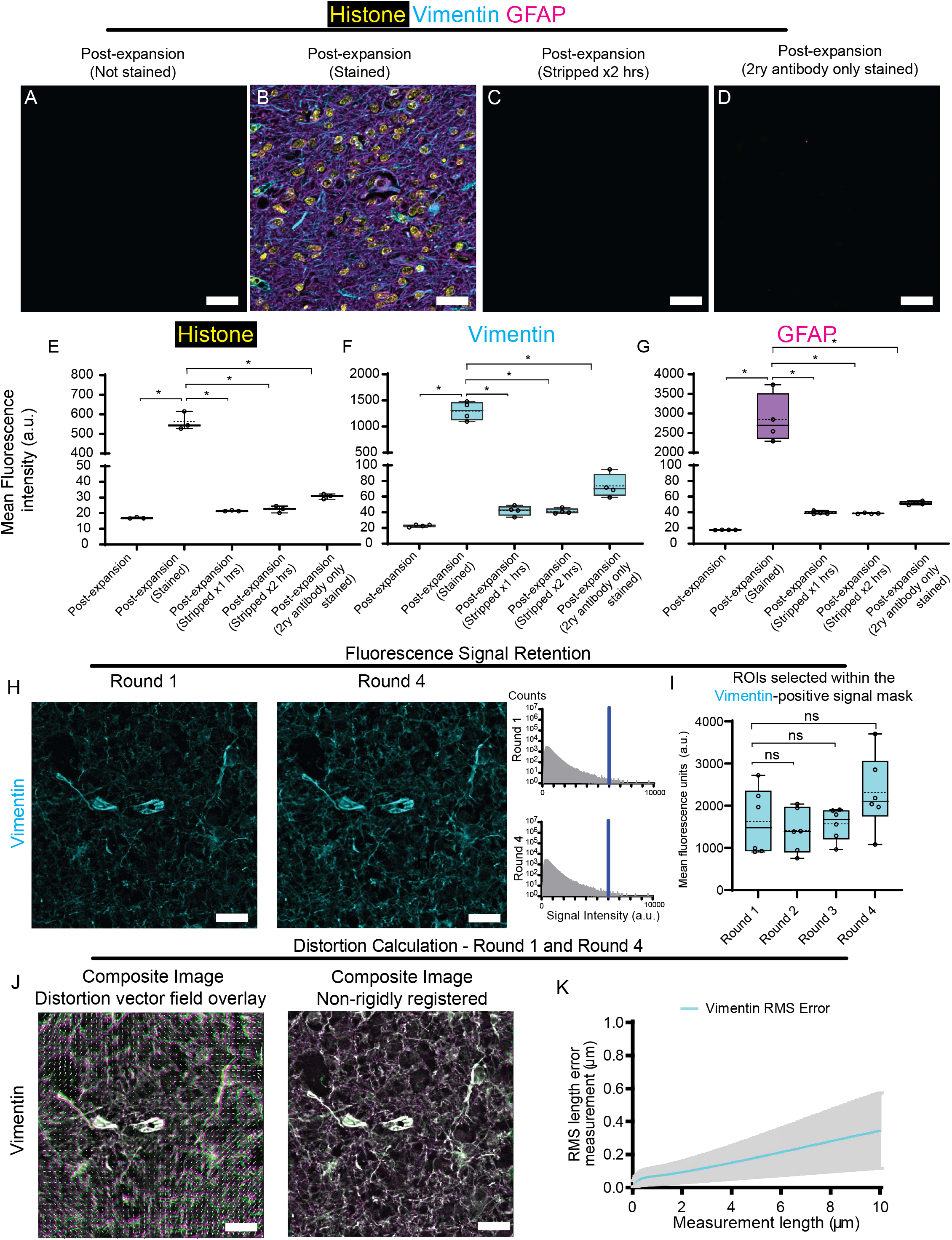
dExPath antibody stripping clears fluorescence signals and enables multiple rounds of post-decrowding immunostaining. (A) Representative (n = 4 samples, from 2 patients with 2 samples per patient) confocal images (sum intensity z-projections of image stacks) of an FFPE, 5-µm-thick tissue slice of human high-grade glioma. Sample underwent format conversion (**Supp. Fig. 1A**; tissue deparaffinization and rehydration), anchoring, gelation (**Supp. Fig. 1B**), softening (**Supp. Fig. 1C**), de-crowding (**Supp. Fig. 1D**), no immunostaining, and confocal imaging at 4x linear expansion (**Supp. Fig. 1E**). (B) Sample used for (A) after immunostaining post-decrowding for histone, vimentin, and GFAP, and imaging at ∼4x linear expansion (**Supp. Fig. 1E**). Sum intensity z-projection of an image stack covering the biological thickness of the original z-projection (used for all expanded images throughout this figure); image was of the same field of view as in (A), using identical hardware settings. (C) Sample used for (A) after antibody stripping for 2 hrs (**Supp. Fig. 1F**), expansion, and imaging at 4x linear expansion; image was of the same field of view as in (A), using identical settings. (D) Sample used for (A) after additional immunostaining with only the fluorescent secondary (abbreviated 2ry throughout this figure) antibodies, using the same staining conditions as used in (B), and confocal imaging at 4x linear expansion (**Supp. Fig. 1E**); image was of the same field of view as in (A), using identical settings. (E-G) Mean fluorescence intensities, from (from left to right in each graph) post-expansion (not stained, as in A), post-expansion (stained, as in B), post-expansion (stripped for 1 hr), post-expansion (stripped for 2 hrs, as in C), and post-expansion (2ry antibody-only stained, as in D), averaged across ROIs that exhibited prominent fluorescence for (E) histone, (F) vimentin, or (G) GFAP in the post-expansion (stained, as in B) state; colors correspond to the colors in (B) (n = 4 samples, from 2 patients with 2 samples per patient). Box plot: individual values (open circles; 2 measurements were acquired from each patient), median (middle line), mean (dotted line), first and third quartiles (lower and upper box boundaries), lower and upper raw values (whiskers). Statistical testing: 2-tailed paired t-test (non-Bonferroni corrected) was applied to post-expansion (stained) vs. all other post-expansion mean fluorescence intensities for each spectral channel. *, p < 0.05. (H) Representative (n = 3 samples, each from a different patient) confocal images of FFPE, 5-µm-thick tissue of human high-grade glioma. The sample underwent format conversion (**Supp. Fig. 1A**; tissue deparaffinization and rehydration), anchoring, gelation (**Supp. Fig. 1B**), softening (**Supp. Fig. 1C**), decrowding (**Supp. Fig. 1D**), and immunostaining post-decrowding for vimentin, and confocal imaging at ∼4x linear expansion (**Supp. Fig. 1E**) after one round (left panel, Round 1) of post-decrowding staining. The sample then underwent three additional sequential rounds of antibody stripping (**Supp. Fig. 1F**), re-staining post-decrowding with anti-vimentin, and 4x linear expansion (**Supp. Fig. 1E**, for a total of four rounds of immunostaining (right panel, Round 4). Shown in both cases is the sum intensity z-projection of the confocal image stack, corresponding to the biological thickness of the z-projection in Round 1, taken under identical settings and of the same field of view as in Round 1. For display purposes, histograms of pixel values for vimentin images were adjusted so that 1% of the pixels were saturated (histograms for Rounds 1 and 4 are shown to the right of the Round 4 image, top and bottom, respectively). Vertical blue line, upper look-up table (LUT) limit (so that 1% of pixels are saturated). (I) Mean fluorescence intensities, from Round 1 to Round 4 post-expansion images (raw image data, not adjusted as in the images of H), averaged across ROIs that exhibited prominent fluorescence for vimentin (n = 3 samples, each from a different patient). Box plot: individual values (open circles; 2 measurements were acquired from each patient), median (middle line), mean (dotted line), first and third quartiles (lower and upper box boundaries), lower and upper raw values (whiskers). Statistical testing: 2-tailed paired t-test (non-Bonferroni corrected) was applied on Round 1 vs each of Round 2 through 4, post-expansion mean fluorescence intensities. Statistical significance: ns, not significant. (J) (Left panel) Same sample as in (H), with composite image overlaying Round 1 (magenta) and Round 4 (green) post-expansion images prior to non-rigid registration. Distortion vector field overlay (white arrows) derived from non-rigid registration. (Right panel) Composite image of Round 1 and 4 as in left panel, following non-rigid registration. (K) RMS length measurement errors obtained by comparing Round 1 and Round 4 post-expansion images such as those of (J) (n = 3 samples, each from a different patient). Line, mean; shaded area, standard deviation. Scale bars (in biological units): (A-D) 90 µm; (H) 30 µm; (J) 30 µm.

**Supp Fig. 8.**
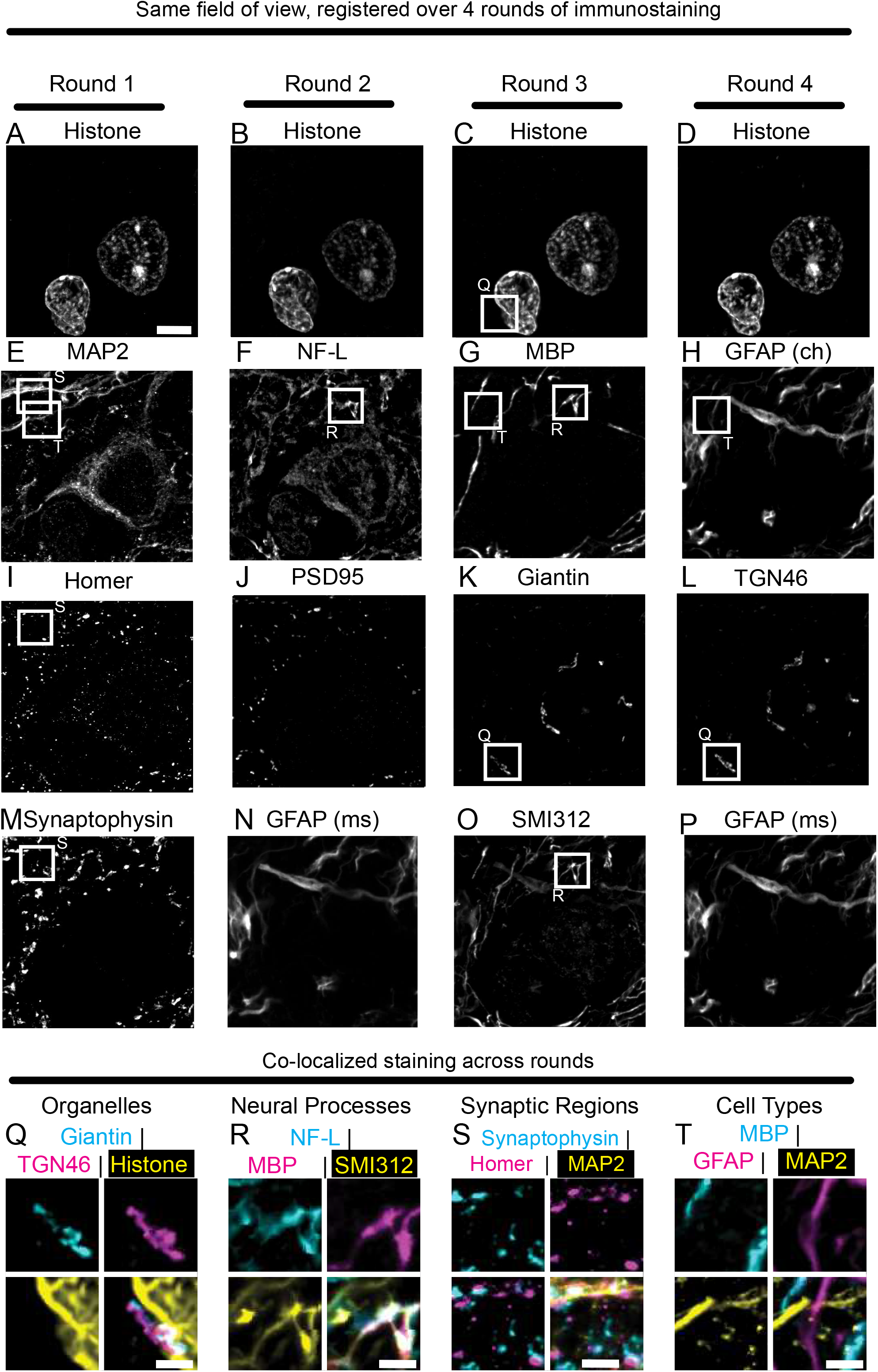
dExPath antibody multiplexing of archival pathology samples of human cortex. (A-L) Example confocal images of the same field of view, for 10 distinct protein targets, from a formalin-fixed paraffin-embedded (FFPE), 5-µm-thick tissue of human brain cortex, which underwent format conversion (as in **Supp. Fig. 1A**; including tissue deparaffinization and rehydration), anchoring, gelation (as in **Supp. Fig. 1B**), softening (as in **Supp. Fig. 1C**), decrowding (as in **Supp. Fig. 1D**), and 4 total rounds of post-decrowding immunostaining (as in **Supp. Fig. 1E**), alternating with antibody stripping treatment (as in **Supp. Fig. 1F**). The protein targets included (A-D) histone, which was a common target across all 4 rounds of staining, to provide a constant landmark for image registration across separate rounds; (E) microtubule-associated protein 2 (MAP2); (F) neurofilament light chain (NF-L); (G) myelin-basic protein (MBP); (H) glial fibrillary acidic protein (GFAP) stained by a polyclonal antibody raised in chicken (“ch”, for contrast to a differently sourced GFAP antibody, below); (I) Homer, a post-synaptic protein; (J) Post-synaptic density 95 (PSD95), another post-synaptic protein; (K) Giantin, a cis-Golgi marker; (L) TGN46, a trans-Golgi marker; (M) synaptophysin, a pre-synaptic protein; (N) GFAP, stained by a mouse monoclonal (“ms”); (O) SMI-312, a monoclonal antibody against phosphorylated neurofilament subunits; (P) GFAP, stained as in N. White boxes are zoomed-in, and overlaid, in Q-T. (Q) Magnified views of the regions inside the solid white boxes in C, K, and L. Upper left, giantin; upper right, TGN46; lower left, histone; lower right, overlay of the other 3 images. (R) Magnified views of the regions inside the solid white boxes in F, G, and O. Upper left, NF-L; upper right, MBP; lower left, SMI-312; lower right, overlay of the other 3 images. (S) Magnified views of the regions inside solid white boxes in E, I, and M. Upper left, synaptophysin; upper right, Homer; lower left, overlay of Homer and synaptophysin; lower right, overlay of Homer, synaptophysin, and MAP2. (T) Magnified views of the regions inside solid white boxes in E, G, and H. Upper left, MBP; upper right, GFAP; lower left, MAP2; lower right, overlay of the other 3 images. All images are sum intensity z-projections of image stacks acquired with confocal microscopy. Brightness and contrast settings: first set by the ImageJ auto-scaling function, and then manually adjusted (by raising the minimum-intensity threshold and lowering the maximum-intensity threshold) to improve contrast for the stained structures of interest. Pixel intensity values were deliberately saturated for a subset of pixels, to facilitate visualizing the spatial distribution of the stains. The adjustments were individually performed for each image. Scale bars (in biological units): (A) 3.5 µm (Panels A-P show the same field of view in the tissue sample). (Q-T) 1.1 µm. Linear expansion factor, 3.9 x.

**Supplementary Table 1.**
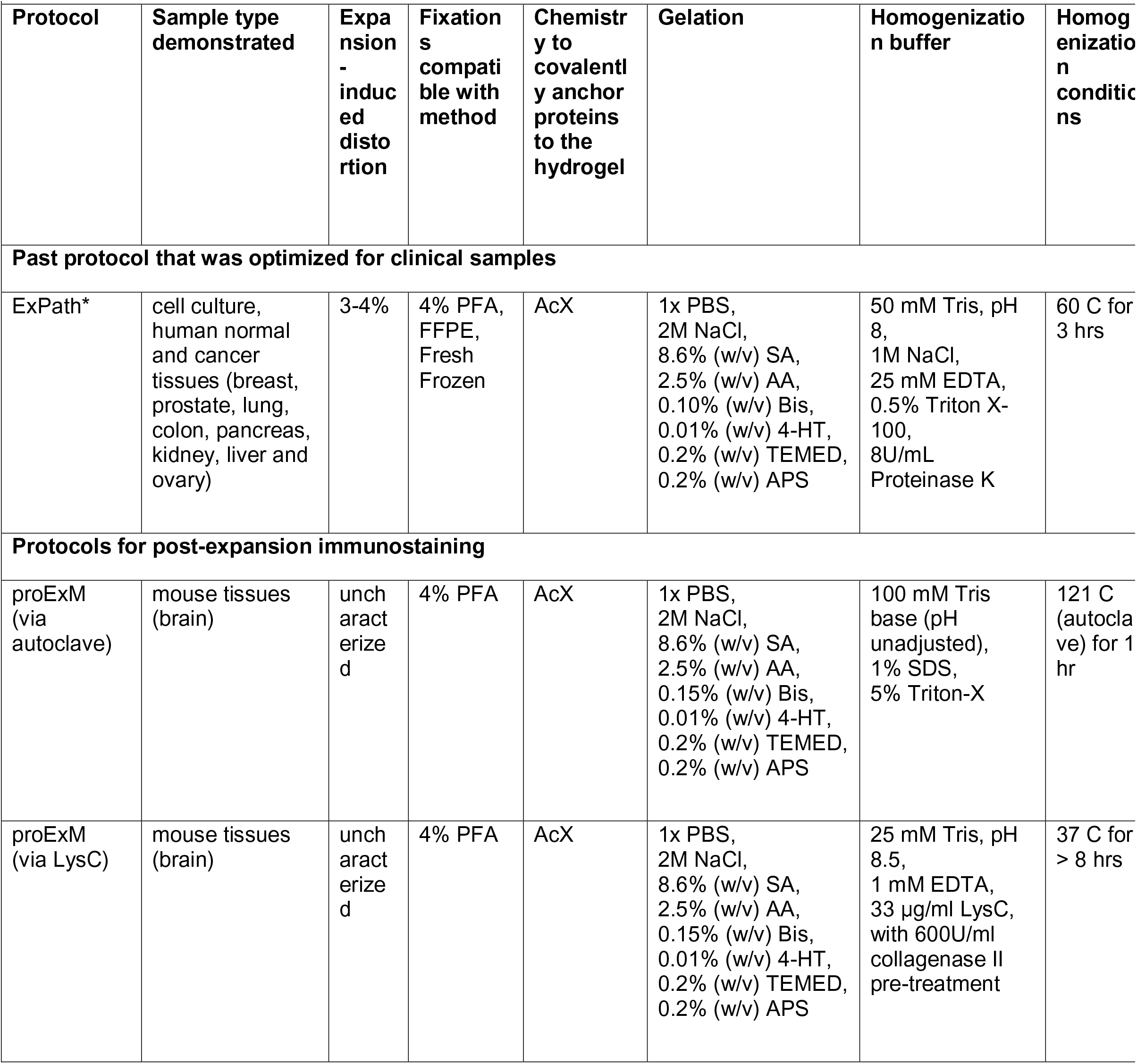
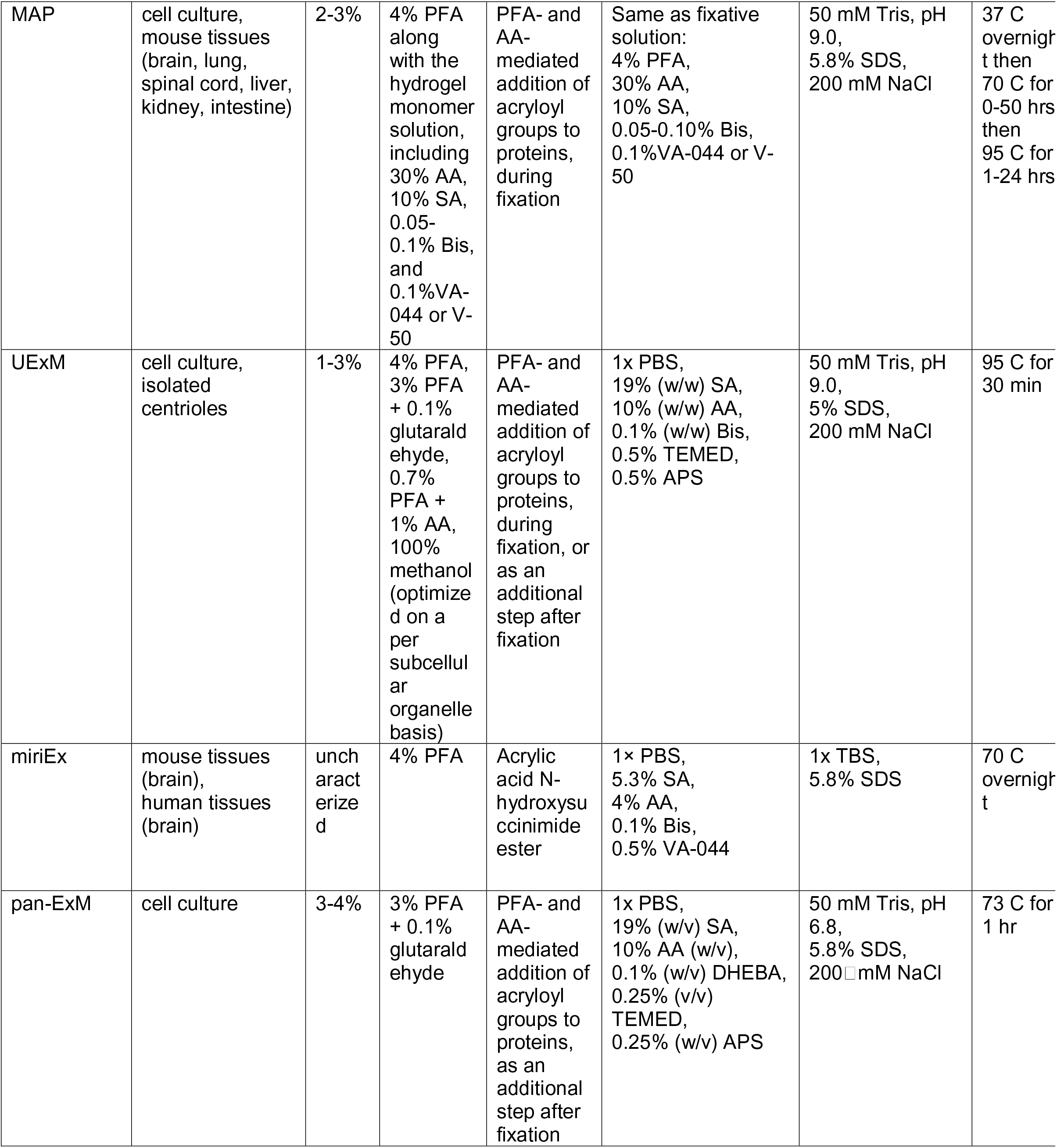
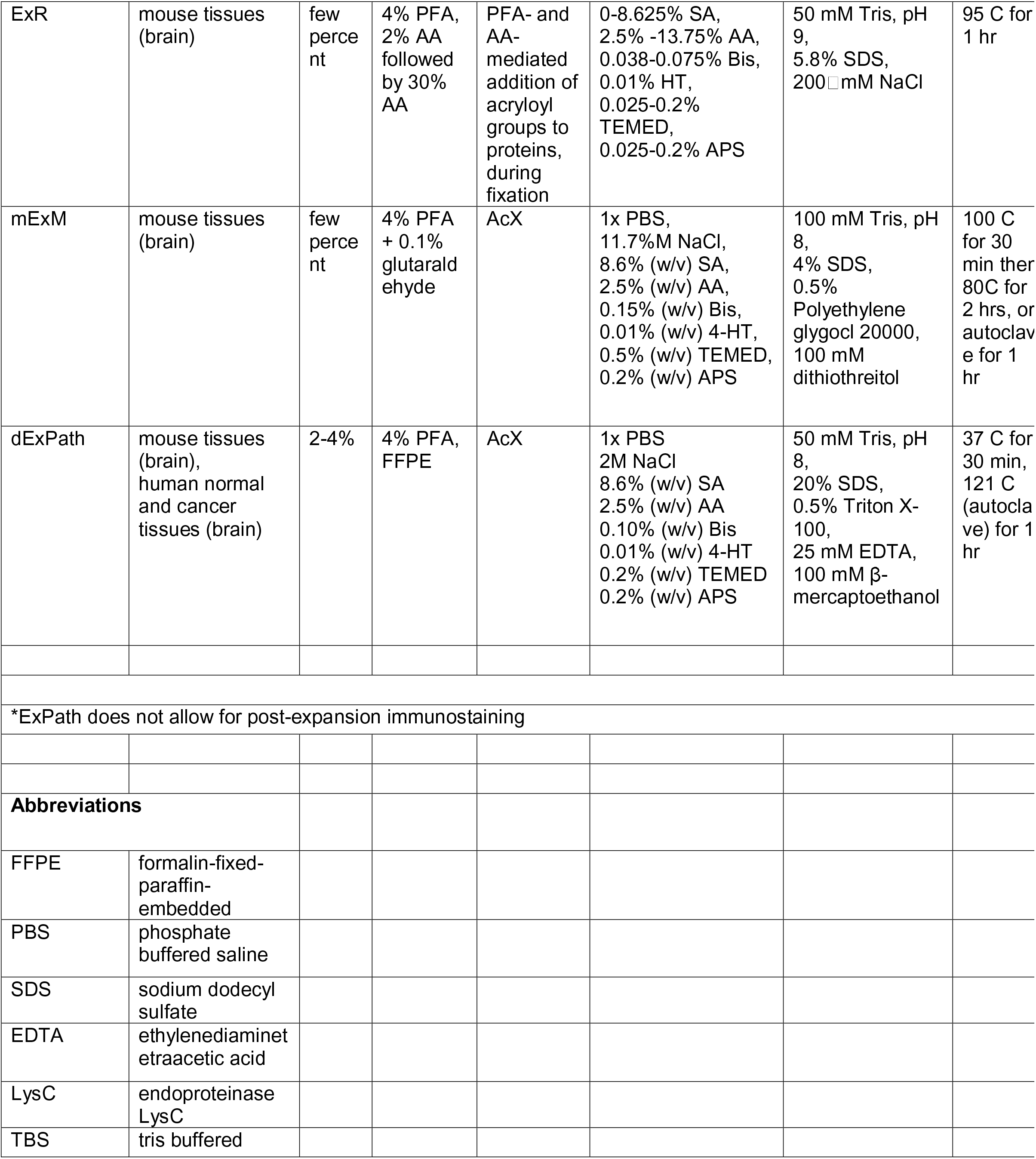
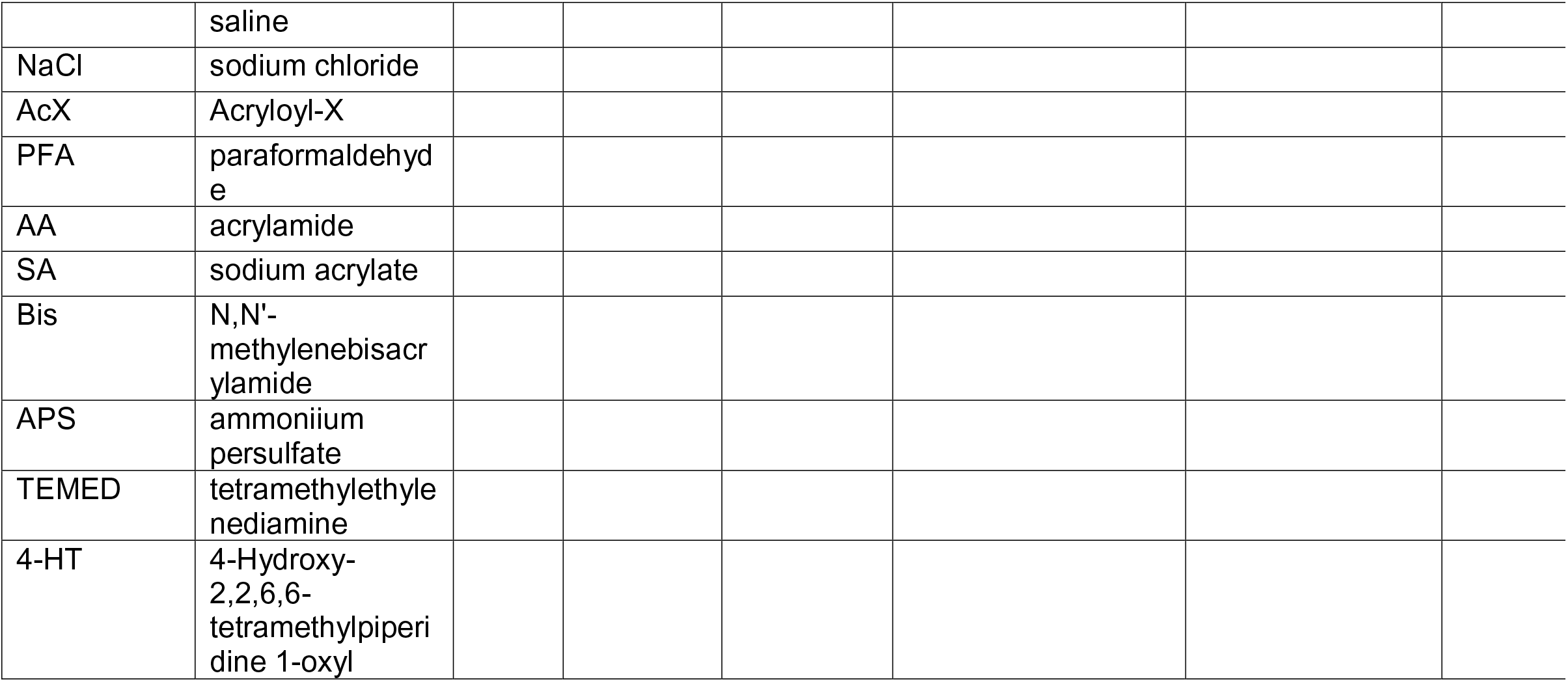
Comparison of tissue expansion protocols

| Supplementary Table 1. Comparison of tissue expansion protocols |  |  |  |  |  |  |  |
| --- | --- | --- | --- | --- | --- | --- | --- |
| Protocol | Sample type demonstrated | Expansion - induced distortion | Fixations compatible with method | Chemistry to covalently anchor proteins to the hydrogel | Gelation | Homogenization buffer | Homogenization conditions |
| **Past protocol that was optimized for clinical samples** |  |  |  |  |  |  |  |
| ExPath* | cell culture, human normal and cancer tissues (breast, prostate, lung, colon, pancreas, kidney, liver and ovary) | 3-4% | 4% PFA, FFPE, Fresh Frozen | AcX | 1x PBS, 2M NaCl, 8.6% (w/v) SA, 2.5% (w/v) AA, 0.10% (w/v) Bis, 0.01% (w/v) 4-HT, 0.2% (w/v) TEMED, 0.2% (w/v) APS | 50 mM Tris, pH 8, 1M NaCl, 25 mM EDTA, 0.5% Triton X-100, 8U/mL Proteinase K | 60 C for 3 hrs |
| **Protocols for post-expansion immunostaining** |  |  |  |  |  |  |  |
| proExM (via autoclave) | mouse tissues (brain) | uncharacterized | 4% PFA | AcX | 1x PBS, 2M NaCl, 8.6% (w/v) SA, 2.5% (w/v) AA, 0.15% (w/v) Bis, 0.01% (w/v) 4-HT, 0.2% (w/v) TEMED, 0.2% (w/v) APS | 100 mM Tris base (pH unadjusted), 1% SDS, 5% Triton-X | 121 C (autoclave) for 1 hr |
| proExM (via LysC) | mouse tissues (brain) | uncharacterized | 4% PFA | AcX | 1x PBS, 2M NaCl, 8.6% (w/v) SA, 2.5% (w/v) AA, 0.15% (w/v) Bis, 0.01% (w/v) 4-HT, 0.2% (w/v) TEMED, 0.2% (w/v) APS | 25 mM Tris, pH 8.5, 1 mM EDTA, 33 µg/ml LysC, with 600U/ml collagenase II pre-treatment | 37 C for > 8 hrs |
| MAP | cell culture, mouse tissues (brain, lung, spinal cord, liver, kidney, intestine) | 2-3% | 4% PFA along with the hydrogel monomer solution, including 30% AA, 10% SA, 0.05-0.1% Bis, and 0.1% VA-044 or V-50 | PFA- and AA-mediated addition of acryloyl groups to proteins, during fixation | Same as fixative solution: 4% PFA, 30% AA, 10% SA, 0.05-0.10% Bis, 0.1% VA-044 or V-50 | 50 mM Tris, pH 9.0, 5.8% SDS, 200 mM NaCl | 37 C overnight then 70 C for 0-50 hrs then 95 C for 1-24 hrs |
| UExM | cell culture, isolated centrioles | 1-3% | 4% PFA, 3% PFA + 0.1% glutaraldehyde, 0.7% PFA + 1% AA, 100% methanol (optimized on a per subcellular organelle basis) | PFA- and AA-mediated addition of acryloyl groups to proteins, during fixation, or as an additional step after fixation | 1x PBS, 19% (w/w) SA, 10% (w/w) AA, 0.1% (w/w) Bis, 0.5% TEMED, 0.5% APS | 50 mM Tris, pH 9.0, 5% SDS, 200 mM NaCl | 95 C for 30 min |
| miriEx | mouse tissues (brain), human tissues (brain) | uncharacterized | 4% PFA | Acrylic acid N-hydroxysuccinimide ester | 1x PBS, 5.3% SA, 4% AA, 0.1% Bis, 0.5% VA-044 | 1x TBS, 5.8% SDS | 70 C overnight |
| pan-ExM | cell culture | 3-4% | 3% PFA + 0.1% glutaraldehyde | PFA- and AA-mediated addition of acryloyl groups to proteins, as an additional step after fixation | 1x PBS, 19% (w/v) SA, 10% AA (w/v), 0.1% (w/v) DHEBA, 0.25% (v/v) TEMED, 0.25% (w/v) APS | 50 mM Tris, pH 6.8, 5.8% SDS, 200 mM NaCl | 73 C for 1 hr |
| ExR | mouse tissues (brain) | few percent | 4% PFA, 2% AA followed by 30% AA | PFA- and AA-mediated addition of acryloyl groups to proteins, during fixation | 0-8.625% SA, 2.5% -13.75% AA, 0.038-0.075% Bis, 0.01% HT, 0.025-0.2% TEMED, 0.025-0.2% APS | 50 mM Tris, pH 9, 5.8% SDS, 200 mM NaCl | 95 C for 1 hr |
| mExM | mouse tissues (brain) | few percent | 4% PFA + 0.1% glutaraldehyde | AcX | 1x PBS, 11.7% NaCl, 8.6% (w/v) SA, 2.5% (w/v) AA, 0.15% (w/v) Bis, 0.01% (w/v) 4-HT, 0.5% (w/v) TEMED, 0.2% (w/v) APS | 100 mM Tris, pH 8, 4% SDS, 0.5% Polyethylene glycol 20000, 100 mM dithiothreitol | 100 C for 30 min then 80C for 2 hrs, or autoclave for 1 hr |
| dExPath | mouse tissues (brain), human normal and cancer tissues (brain) | 2-4% | 4% PFA, FFPE | AcX | 1x PBS 2M NaCl 8.6% (w/v) SA 2.5% (w/v) AA 0.10% (w/v) Bis 0.01% (w/v) 4-HT 0.2% (w/v) TEMED 0.2% (w/v) APS | 50 mM Tris, pH 8, 20% SDS, 0.5% Triton X-100, 25 mM EDTA, 100 mM β-mercaptoethanol | 37 C for 30 min, 121 C (autoclave) for 1 hr |

\*ExPath does not allow for post-expansion immunostaining

|  |  |
| --- | --- |
| **Abbreviations** |  |
| FFPE | formalin-fixed-paraffin-embedded |
| PBS | phosphate buffered saline |
| SDS | sodium dodecyl sulfate |
| EDTA | ethylenediaminetetraacetic acid |
| LysC | endoprotease LysC |
| TBS | tris buffered saline |
| NaCl | sodium chloride |
| AcX | Acryloyl-X |
| PFA | paraformaldehyde |
| AA | acrylamide |
| SA | sodium acrylate |
| Bis | N,N'-methylenebisacrylamide |
| APS | ammonium persulfate |
| TEMED | tetramethylethylenediamine |
| 4-HT | 4-Hydroxy-2,2,6,6-tetramethylpiperidine 1-oxyl |

**Supplementary Table 2.** Antibodies Used

| **Supplementary Table 2. Antibodies Used** |  |  |  |
| --- | --- | --- | --- |
| **Primary Antibodies** |  |  |  |
| **Target** | **Host** | **Manufacturer** | **Catalog No.** |
| α-SMA (1A4) | Mouse | Agilent | M085129-2 |
| Bassoon (SAP7F407) | Mouse | Abcam | ab82958 |
| Collagen IV | Goat | Millipore Sigma | AB769 |
| Collagen IV | Rabbit | Abcam | ab6586 |
| GFAP (GA5) | Mouse | ThermoFisher Scientific | 14-9892-82 |
| GFAP | Chicken | Novus Biologicals | NBP105198 |
| GFAP | Chicken | Aves Labs | GFAP |
| Giantin | Rabbit | Biolegend | 924302 |
| Iba1 | Rabbit | WAKO | 019-19741 |
| Histone-H3 | Goat | Abcam | ab1791 |
| Homer 1 | Rabbit | Synaptic Systems | 160 003 |
| Laminin | Chicken | LSBio | LS-C96142 |
| MAP2 | chicken | Novus Biologicals | NB300213 |
| MBP | Chicken | Millipore Sigma | AB9348 |
| NEFL (NF70) | Chicken | ThermoFisher Scientific | PA1-10000 |
| Neurofilament-M | Rabbit | Millipore Sigma | AB1987 |
| PSD95 | Rabbit | Cell Signaling Technology | 2507 |
| SMI312 | Mouse | Biolegend | 837904 |
| Synaptophysin (SVP-38) | Mouse | Millipore Sigma | S5768 |
| Vimentin | Chicken | Abcam | ab24525 |
| ZO-1 (ZO1-1A12) | Mouse | ThermoFisher Scientific | 33-9100 |
| **Secondary Antibodies** |  |  |  |
| **Target** | **Host** | **Manufacturer** | **Catalog No.** |
| AlexaFluor 350, anti-goat | Donkey | ThermoFisher Scientific | A10040 |
| AlexaFluor 488, anti-chicken | Donkey | Jackson ImmunoResearch Laboratories | 703545155 |
| AlexaFluor 488, anti-chicken | Goat | ThermoFisher Scientific | A11039 |
| AlexaFluor 546, anti-rabbit | Donkey | ThermoFisher Scientific | A10040 |
| AlexaFluor 546, anti-rabbit | Goat | ThermoFisher Scientific | A10035 |
| CF 640, anti-mouse | Donkey | Biotium | 20177 |

